# Impact of age and surface irregularities on intersegmental and inter-joint coordination during gait

**DOI:** 10.64898/2026.05.22.727144

**Authors:** Antoine Cohic, Cloé Dussault-Picard, Silvère De Freitas, Yosra Cherni

## Abstract

Uneven walking surfaces require adjustments in motor strategies and can thus provide insights into the neuromuscular changes underlying maturation. Also, coordination metrics and variability offer a richer description of motor control mechanisms than standard spatiotemporal parameters, constituting a more sensitive approach to characterize developmental changes. This study aimed to investigate the effects of uneven surfaces on intersegmental coordination, as well as inter-joint coordination and its variability, and to assess the differences in adaptation between age groups when walking on uneven surfaces.

Seventy participants (2-29 years), divided into four age groups, completed gait trials on an even and two levels of uneven surfaces while equipped with reflective markers. Mean absolute relative phase and deviation phase of the knee-hip and ankle-knee joint pairs were computed to characterize lower limb inter-joint coordination and variability. In addition, the organization and density of whole-body intersegmental coordination were assessed using correlation networks built from marker acceleration data.

Uneven surfaces induced more in-phase inter-joint coupling, reduced network density and increased variability across all age groups. While the organization of intersegmental coordination remained stable, older participants exhibited denser networks, reflecting refined segmental interactions. In contrast, younger participants showed more in-phase joint coordination and higher variability suggesting less mature motor control. The age-related inter-joint coordination differences were emphasized on uneven surfaces, likely reflecting the maturation-related ability to modulate spinal locomotor patterns via supraspinal control, thereby increasing adaptation to environmental perturbations.

**Highlights:**

– Uneven surfaces induce more in-phase inter-joint coordination.
– Uneven surfaces accentuate differences in locomotor strategies across development.
– Kinectome density may be a promising indicator of locomotor maturation.
– Coordinative variability decreased with neuromotor development.

## 1. Introduction

Human gait undergoes progressive maturation throughout childhood and adolescence, reflecting the development of neuromuscular control, coordination, and balance (Froehle et al., 2013). These developmental changes are particularly evident when individuals are exposed to challenging environments during walking, such as irregular surfaces (Gentle et al., 2016), which require continuous sensorimotor adjustments, coordinated movements and dynamic balance control (Dussault-Picard, Mohammadyari, et al., 2022). Moreover, in clinical populations such as children with cerebral palsy, walking on irregular surfaces can reveal motor impairments that are not apparent on regular surfaces (Dussault-Picard et al., 2023). Under such conditions, walkers commonly adopt compensatory strategies including reduced speed, increased base of support, and greater spatiotemporal variability (Brognara et al., 2025; Dussault-Picard et al., 2025), reflecting attempts to maintain stability and adapt to environmental perturbations.

Much of the current literature characterize gait adaptations using spatiotemporal parameters such as stride length, cadence, and support times (Dussault-Picard, Mohammadyari, et al., 2022; Inns et al., 2025). While these metrics summarize global walking strategies, they provide a limited and reductive understanding of the motor control mechanisms and coordination strategies that emerge throughout maturation in challenging locomotor contexts (Brognara et al., 2025; Dussault-Picard, Mohammadyari, et al., 2022). Indeed, similar spatiotemporal profiles may conceal distinct neuromechanical strategies, highlighting the need for more sensitive and multidimensional analytical approaches (Brognara et al., 2025).

Growing evidence suggests that multidimensional parameters, including coordination metrics and their variability, are sensitive to developmental differences (Goetschalckx et al., 2024) and motor control deficits (Dussault-Picard, Ippersiel, et al., 2022; Dussault-Picard et al., 2023), providing information beyond those captured by monoarticular kinematic measures. In this context, intersegmental and inter-joint coordination analyses have been used to investigate mechanisms by which the central nervous system organizes movement across body segments to achieve stable and efficient locomotion (Carollo et al., 2018; Ippersiel et al., 2021, 2022; Stergiou et al., 2001; van Emmerik et al., 2014). Thus, coordination analysis and its variability may offer a more comprehensive perspective on motor strategies and the mechanisms underlying adaptive gait behaviour during maturation.

Existing quantitative approaches to assess inter-joint or intersegmental coordination during gait include kinematic analysis methods such as vector coding (Celestino et al., 2019; Heiderscheit et al., 2002; Needham et al., 2014) and phase-based techniques grounded in dynamical systems theory (Burgess-Limerick et al., 1993; Hamill et al., 1999; Kelso, 1995; Stergiou & Decker, 2011). Among these, continuous relative phase (CRP) has emerged as a widely used method as it provides a continuous description of coordination patterns, enabling the identification of in-phase and out-of-phase relationships between joints or segments during the gait cycle (Dussault-Picard et al., 2023; Ippersiel et al., 2022). However, CRP is limited to pairwise analyses and thus offers only a partial representation of coordination at the whole-body level during gait. Complementary approaches capable of capturing the global structure of intersegmental interactions are therefore needed to extend this framework.

The kinectome has recently been proposed as a multidimensional approach to represent whole-body coordination patterns by modelling the interactions among multiple segments within a fully connected network (Troisi Lopez et al., 2022). Therefore, combining CRP and kinectome analyses offers complementary insights, enabling both a detailed characterization of lower limb coordination during gait and a comprehensive representation of whole-body coordination.

This study aims to **1)** assess adaptations in intersegmental and inter-joint coordination of children, adolescents and adults when walking across different levels of uneven surfaces, to **2**) quantify inter-joint coordination variability, and to **3)** investigate the maturation-related evolution of segmental and joint coordination, and their variability during walking. We hypothesized that **1)** uneven surfaces will alter coordination, leading to a more in-phase inter-joint coupling and a less dense intersegmental correlation network, **2)** uneven surfaces will increase inter-joint coordination variability across all age groups, and **3)** coordination patterns and variability will differ between age groups, with lower variability, more out-of-phase coupling and denser correlation networks expected in older participants, reflecting maturation-related differences.

## 2. Methods

### 2.1. Participants

Participants aged 2 to 35 years were recruited. Individuals with a history of musculoskeletal, neurological, cognitive, or psychiatric disorders were excluded. Participants were categorized into four age groups, namely adults (18-35 years), adolescents (12-17 years), children (6-11 years), and young children (2-5 years) (Lally & Valentine-French, 2017; Meyns et al., 2020). All participants or their parents provided informed consent, and all procedures were reviewed and approved by the Research Ethics Board of the Sainte-Justine university hospital (2026-9359).

### 2.2. Data collection

Data acquisition was carried out in the gait analysis laboratory of the rehabilitation centre of the Sainte Justine University Hospital. Participants were instructed to walk at self-selected speed with their usual shoes on three different types of surfaces: even, moderately uneven and highly uneven. The uneven surfaces were made up of polyurethane shock-absorbing floor panels (Terrasensa, Kassel, Germany) on a length of 4 meters, with a maximum vertical variation of 2 cm for the moderately uneven and 5 cm for the highly uneven surface. Each trial extended 2m beyond the start and end of the surfaces to prevent acceleration and deceleration within the measurement area. The sequence of trials was randomized for each participant. Young children performed 5 gait trials on each surface while the other three groups completed 10 gait trials. This choice was made to limit fatigue in young children and did not result in a substantial imbalance in the number of gait cycles between groups, as young children take more steps over the same distance. Participants were equipped with 43 reflective markers placed according to the Plug-in Gait model, and their trajectories were recorded with a 12-camera motion capture system (Vicon Motion Systems Ltd., Oxford, UK) at a sampling frequency of 100 Hz.

### 2.3. Data processing

Labelling and gap filling of marker data, as well as joint angles calculations, were performed using Vicon Nexus (v. 2.14, Vicon Motion Systems Ltd., Oxford, UK), and gait events were manually identified by the same experienced assessor. Data were then exported to Matlab (v.R2025B, MathWorks Inc., Natick, USA) for further processing with custom code.

#### 2.3.1. Intersegmental coordination

Intersegmental coordination was assessed by computing kinectomes, which are symmetrical networks where nodes are represented by markers, and the weighted edges between nodes correspond to the correlations between the linear accelerations of each pair of markers along the anteroposterior axis (Troisi Lopez et al., 2022). To do so, marker trajectories were low-pass filtered (bidirectional Butterworth, 4^th^ order, 6-Hz cut-off) and subsequently differentiated twice to compute accelerations. To reduce redundancy, a subset of 21 markers was selected for analysis, with the retained markers presented in Figure 1. Head markers were combined by calculating the mean acceleration across the four markers. The trunk was represented by C7, T10, LASI, and RASI; the upper limbs by LSHO, LELB, LFIN, RSHO, RELB, and RFIN; and the lower limbs by LTHI, LKNE, LTIB, LANK, LTOE, RTHI, RKNE, RTIB, RANK, and RTOE, with L and R indicating left and right sides, respectively. Marker abbreviations follow the Plug-in Gait model convention.

**Figure 1:**
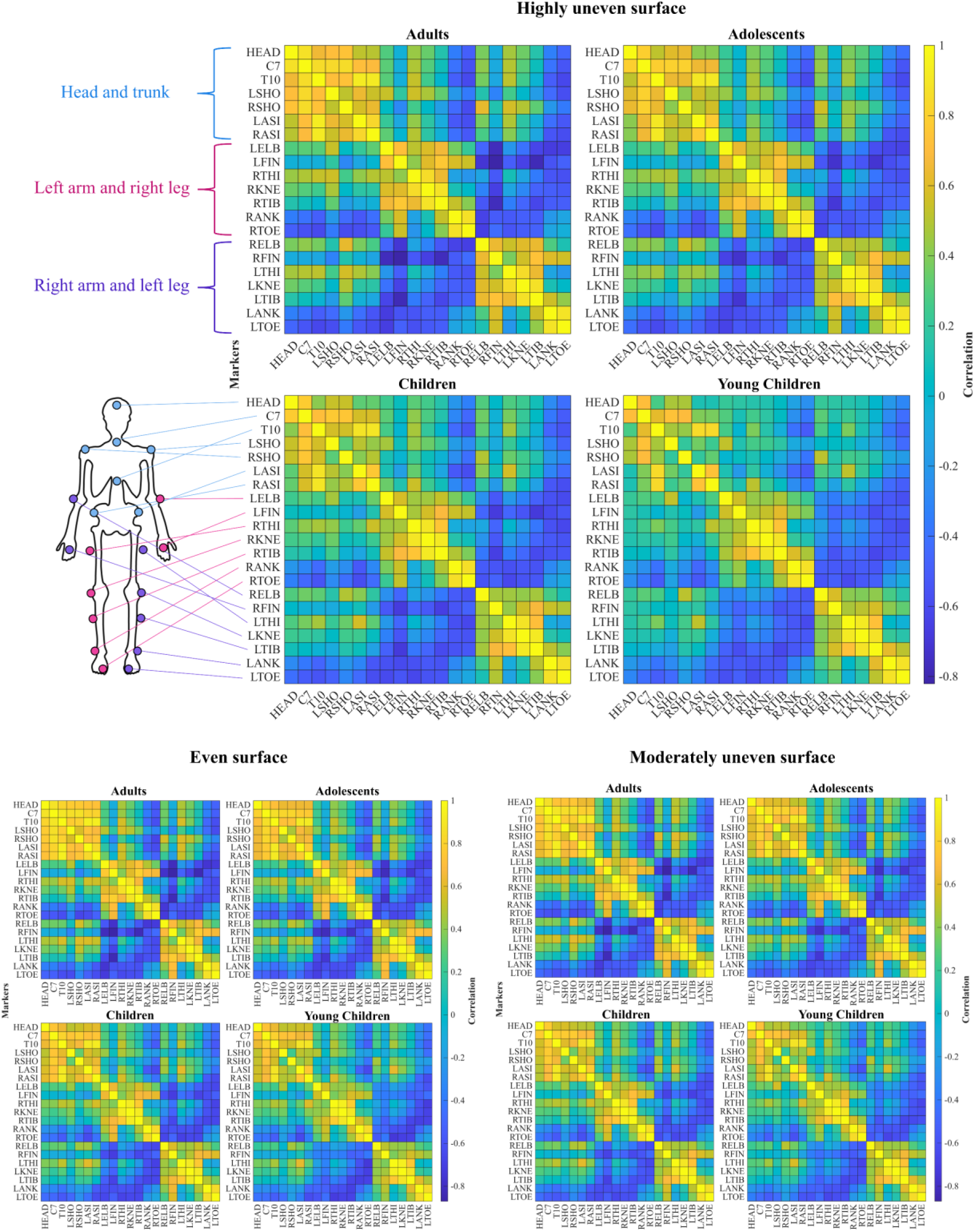
Mean kinectomes across age groups and surfaces

Gait trials were segmented into bilateral cycles, each comprising a gait cycle from both sides, and Pearson’s correlation coefficients were computed between the linear accelerations of all marker pairs along the anteroposterior axis and over each cycle (Troisi Lopez et al., 2022). A mean kinectome was then obtained for each participant and surface by averaging the kinectomes from all bilateral cycles. Finally, mean kinectomes were calculated for each age group and surface.

The Louvain method for community detection (Blondel et al., 2008) was used to organize these mean kinectomes into modules within which markers exhibit similar accelerations during gait. This method was implemented using the Brain Connectivity Toolbox (Rubinov et al., 2009), and the modularity Q used by the algorithm to identify the optimal partition was expressed as in equation (1), following the reformulation proposed by Gómez et al for networks of correlated data (Gómez et al., 2009):

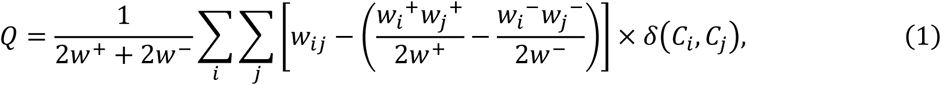

where *w*_*ij*_ is the weight of the edge between nodes *i* and *j*, *w*_*i*_^+^ and *w*_*i*_^−^ are the positive and negative strengths of node *i* (i.e., the sum of the positive or negative weights between *i* and all other nodes), *w*^+^and *w*^−^ are the positive and negative total strengths of the network (i.e., the sum of the positive or negative strength of all nodes) and δ(*C*_*i*_, *C*_*j*_) is the Kronecker delta function between the communities of nodes *i* and *j* (i.e., 1 if nodes *i* and *j* are in the same community, 0 otherwise).

Whole-body intersegmental coordination was assessed by calculating the density D of the marker network, defined as the mean of the absolute weights between nodes, and expressed as in equation (2):

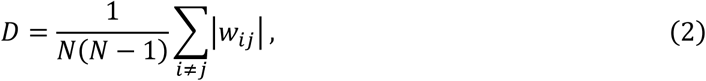

where N is the number of nodes.

#### 2.3.2. Inter-joint coordination and its variability

Inter-joint coordination was assessed using the CRP method, obtained by calculating the phase angles difference between two joints. This method was performed on the lower limbs, for the knee-hip and ankle–knee joint pairs in the sagittal plane (i.e., flexion/extension) (Dussault-Picard et al., 2023).

To this end, joint kinematic signals from all gait trials were first low-pass filtered (bidirectional Butterworth, 4^th^ order, 6-Hz cut-off), segmented into cycles (consecutive heel strikes from the same foot), and padded with spline extrapolation over 10 points on both ends to avoid edge effects (Ippersiel et al., 2019). Signals were then amplitude-centered around zero and phase angles were computed using the Hilbert transform method (Lamb & Stöckl, 2014). Padding points were subsequently removed, and CRP was calculated for each cycle as the absolute difference between the phase angles of corresponding joint pairs. To address potential discontinuities across the gait cycle, CRP values above 180 were transformed by subtracting them from 360, thus mapping all values to a 0–180° range (Ippersiel et al., 2021). A value of 0° indicates that the joints are moving fully in-phase, whereas a value of 180° reflects fully out-of-phase coupling (Burgess-Limerick et al., 1993). Finally, CRP curves were time-normalized to 101 points. Full details of the CRP calculations are available in the Supplementary Material.

Mean absolute relative phase (MARP) (i.e., the mean of the CRP curves) and the deviation phase (DP) (i.e., the standard deviation of the CRP curves), representing the variability of inter-joint coordination, were calculated for each participant and surface over all cycles of both legs.

### 2.4. Statistical analyses

To assess the main and interaction effects of age and surface, a 2-way mixed analysis of variance (ANOVA) was performed for each of our three metrics (Density, MARP and DP).

For density, normality of residuals was assessed by visual inspection of Q-Q plots. If significant main or interaction effects were found, parametric (paired or unpaired t-tests) or non-parametric (Wilcoxon signed-rank or rank-sum tests) post hoc tests with Bonferroni correction (Dunn, 1961) were applied depending on whether the normality assumption was met.

For the continuous MARP and DP metrics, statistical analyses were conducted using the one-dimensional Statistical Parametric Mapping (SPM) software (spm1D, v.M.0.4.52) (Pataky, 2010). Normality of the residual fields was assessed using D’Agostino’s K^2^ test, and post hoc tests were performed using parametric SPM or non-parametric permutation inference with Bonferroni correction (Pataky et al., 2015). Cluster-level p-values were reported, with clusters defined as contiguous regions of the curves in which the test statistic exceeded a predefined threshold. Cohen’s d values were computed for each point within each cluster, and their mean was taken as the corresponding cluster-level effect size (Dussault-Picard et al., 2023). Cohen’s d values < 0.2 indicate negligible effects, 0.2–0.5 small effects, 0.5–0.8 medium effects, and ≥ 0.8 large effects (Cohen, 1988). Only clusters lasting at least 5% of the gait cycle were discussed (Dussault-Picard et al., 2023).

## 3. Results

Descriptive statistics for participant characteristics are reported as mean (standard deviation). Demographic and anthropometric characteristics of each subgroup are presented in Table 1. On average, adults completed 35.7 (7.1) cycles per surface, while adolescents completed 39.6 (6.1), children 48.0 (10.8) and young children 40.1 (18.1).

**Table 1:** Participant demographic and anthropometric characteristics (Mean ± standard deviation)

| Group | Age (years) | Height (cm) | Weight (kg) | F/M |
| --- | --- | --- | --- | --- |
| Young children | 4.6 (1.1) | 105.1 (10.5) | 17.4 (2.5) | 9/6 |
| Children | 8.6 (1.8) | 133.6 (14.8) | 31.1 (10.8) | 14/5 |
| Adolescents | 14.0 (1.3) | 166.0 (9.8) | 54.1 (12.3) | 11/5 |
| Adults | 24.6 (2.8) | 174.3 (8.6) | 70.4 (11.9) | 11/9 |

### 3.1. Intersegmental coordination based on kinectome analysis

The three marker communities identified by the Louvain method were identical across the four age groups, regardless of the surface, and are presented in Figure 1. These three communities have been classified into functional body segments: the head and trunk, and the two pairs of contralateral upper and lower limbs.

The 2-way mixed ANOVA performed on the density of the networks showed a significant main effect of age (p<0.001) and surface (p<0.001), but no significant group x surface interaction (p=0.776). Visual inspection of the Q-Q plots confirmed that the assumptions underlying the ANOVA were met (Supplementary Figure S1). The results of the parametric post hoc tests are reported in Table 2.

**Table 2:**
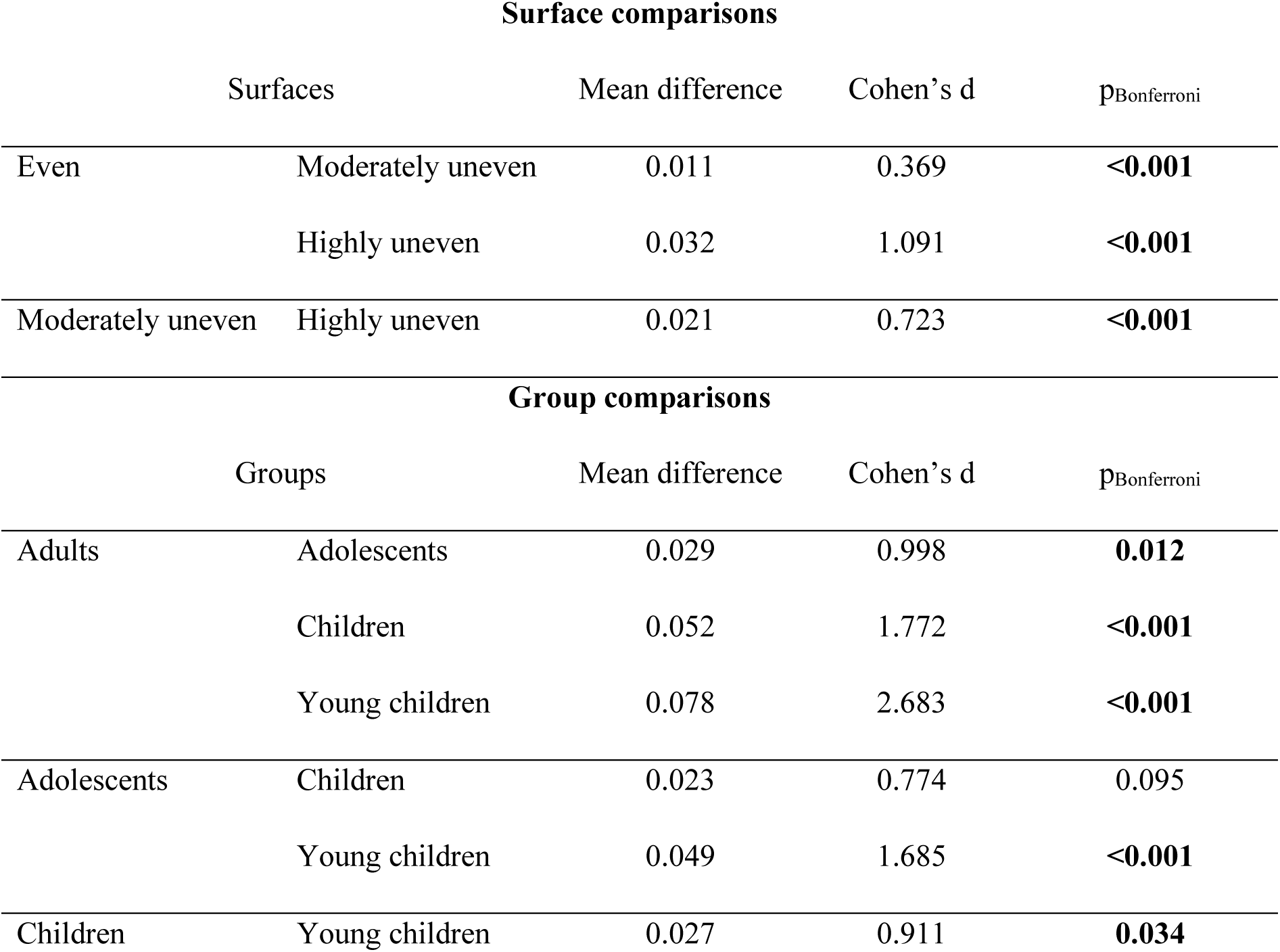
Post hoc surface and group comparisons for density. Bold p-values indicate statistical significance (p<0.05)

When comparing density across surfaces, significant differences were found in all comparisons (Table 2), with the most uneven surface consistently showing lower density. Regarding age group comparisons, a hierarchical organisation across age groups was observed, with the youngest participants exhibiting the lowest mean values under all three surfaces (Figure 2). Significant differences were found between adults and other groups (Adolescents: p=0.008, d=0.998; Children: p<0.001, d=1.772; Young children: p<0.001, d=2.683), between adolescents and young children (p<0.001, d=1.685) and between children and young children (p=0.034, d=0.911).

**Figure 2:**
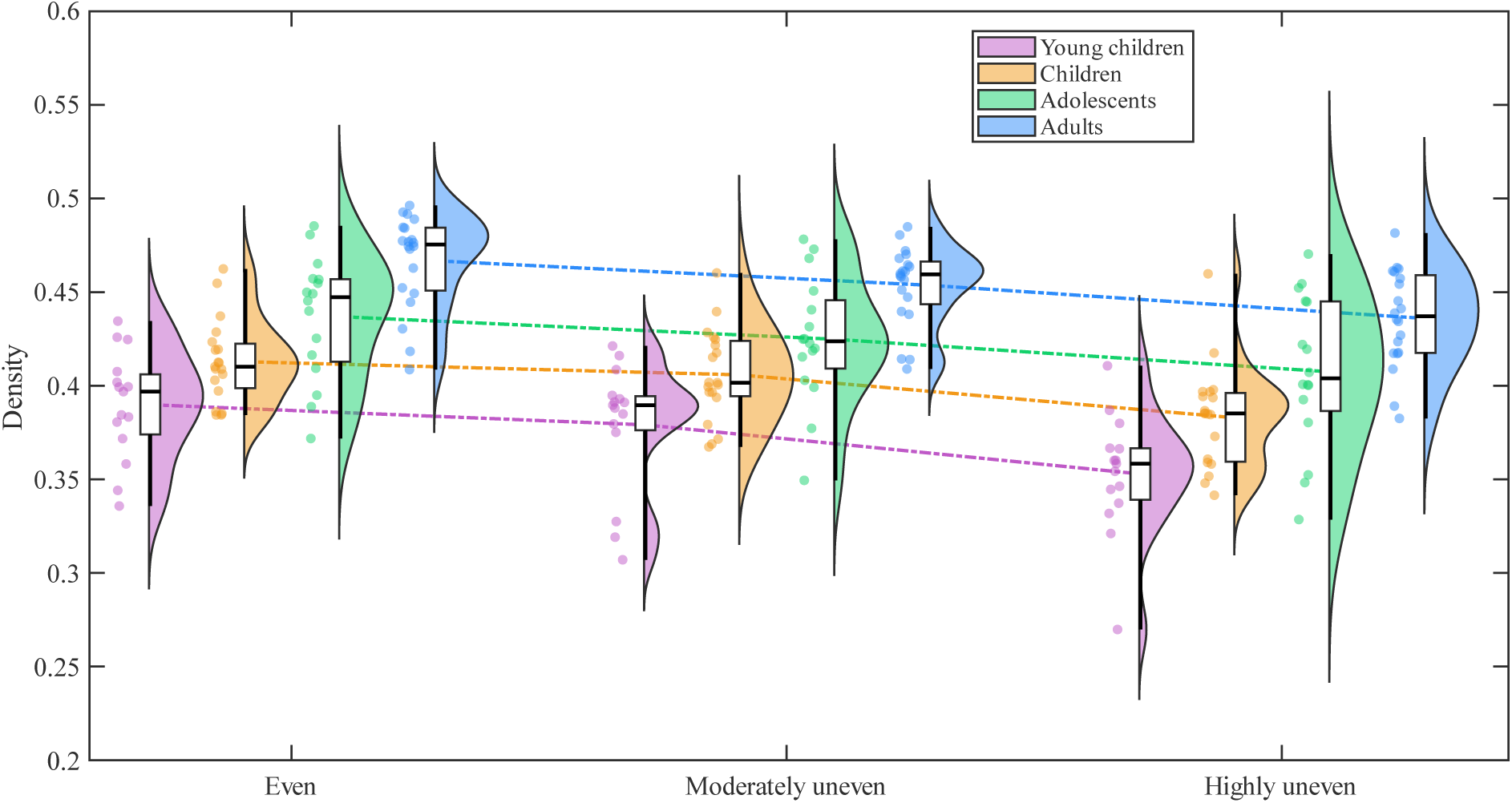
Raincloud plots of density across age groups and surfaces. The dashed lines connect the means of the distributions.

### 3.2. Inter-joint coordination based on CRP analysis

Non-parametric ANOVAs conducted on MARP and DP values for the knee-hip and ankle-knee joint pairs revealed significant main effects of age and surface, as well as significant age x surface interaction, at specific phases of the gait cycle. SPM plots detailing the main effects and interaction for each joint pair and metric are presented in the supplementary material. The results of the subsequent non-parametric post hoc tests are presented in Figures 3 to 10.

**Figure 3:**
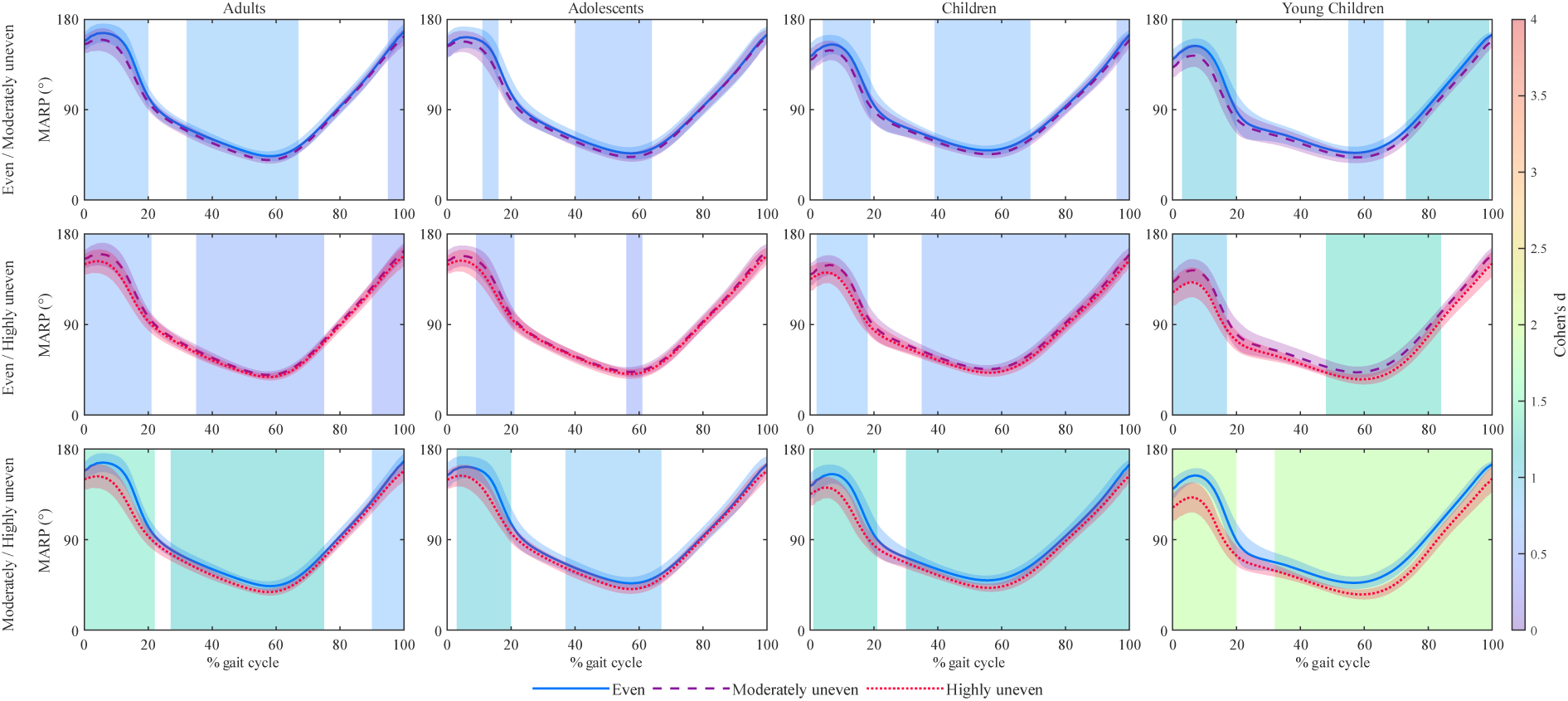
Surface comparisons of MARP curves for the Knee-Hip joint pair. Shaded vertical bands represent areas with a significant difference between curves and are coloured with the associated Cohen’s d effect size

#### 3.2.1. Knee-Hip coordination

##### 3.2.1.1. Mean absolute relative phase

When comparing the highly uneven surface to the even surface (Figure 3), all groups showed a more in-phase knee-hip coordination (i.e., lower MARP) during initial contact and loading response (Adults: 0-22%, p<0.001, d=1.398 – large effect; Adolescents: 3-20%, p<0.001, d=1.099 – large effect; Children: 1-21%, p<0.001, d=1.281 – large effect; Young children: 0-20%, p<0.001, d=1.881 – large effect) and during terminal stance and swing (Adults: 27-75%, p<0.001, d=1.195 – large effect; 90-100%, p<0.001, d=0.778 – medium effect; Adolescents: 37-67%, p<0.001, d=0.857 – large effect; Children: 30-100%, p<0.001, d=1.155 – large effect; Young children: 32-100%, p<0.001, d=1.852 – large effect).

Similar trends were observed when comparing the moderately uneven surface to the even surface during loading response (Adults: 0-20%, p<0.001, d=0.770 – medium effect; Adolescents: 11-16%, p<0.001, d=0.727 – medium effect; Children: 4-19%, p<0.001, d=0.717 – medium effect; Young Children: 3-20%, p<0.001, d=1.048 – large effect), terminal stance and initial swing (Adults: 32-67%, p<0.001, d=0.832 - large effect; Adolescents: 40-64%, p<0.001, d=0.665 – medium effect; Children: 39-69%, p<0.001, d=0.749 – medium effect; Young children: 55-66%, p=0.001, d=0.810 – large effect) in all groups, and during terminal swing in adults (95-100%, p<0.001, d=0.502 – medium effect) and young children (73-99%, p<0.001, d=1.104 – large effect).

Between the moderately and highly uneven surfaces, more in-phase knee-hip coordination was observed during loading response in all groups (Adults: 0-21%, p<0.001, d=0.691 – medium effect; Adolescents: 9-21%, p<0.001, d=0.570 – medium effect; Children: 2-18%, p<0.001, d=0.768 – medium effect; Young children: 0-17%, p<0.001, d=0.942 – medium effect), during terminal stance in adolescents (56-61%, p=0.004, d=0.448 – small effect), and during terminal stance and swing in young children (48-84%, p<0.001, d=1.308 – large effect), children (35-100%, p<0.001, d=0.646 – medium effect) and adults (35-75%, p<0.001, d=0.544 – medium effect; 90-100%, p<0.001, d=0.435 – small effect) when walking on the highly uneven surface.

Regarding age group comparisons (Figure 4), young children showed a more in-phase coordination than the other groups and this tendency was observed over a broader portion of the gait cycle on uneven surfaces. Indeed, when compared to adults, the more in-phase pattern of young children occurred only at initial contact on the even surface (0-9%, p<0.001, d=1.657 – large effect), but extended to the loading phase on the moderately uneven surface (0-8%, p<0.001, d=1.726 – large effect; 13-25%, p<0.001, d=1.519 – large effect), and to the initial swing on the highly uneven surface (0-33%, p<0.001, d=1.743 – large effect; 65-82%, p<0.001, d=1.761 – large effect).

**Figure 4:**
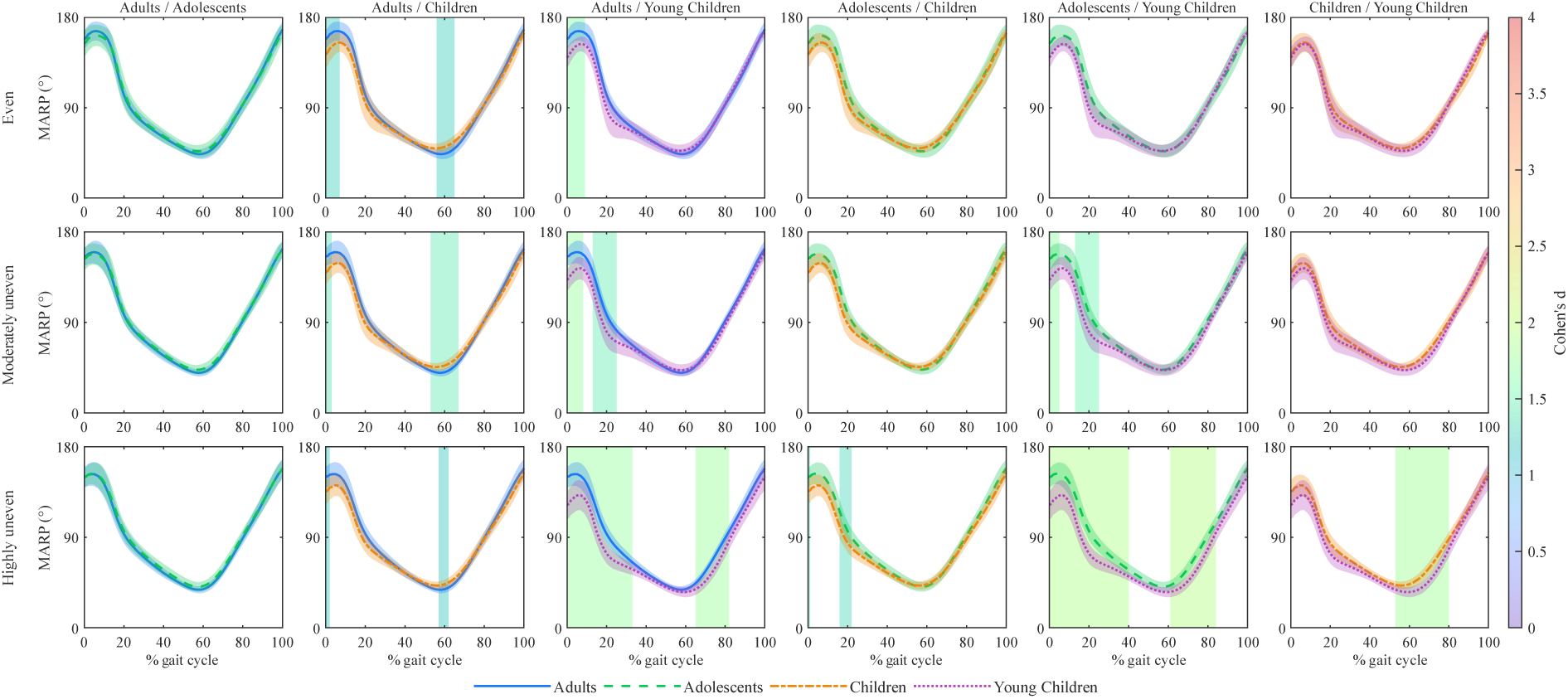
Group comparisons of MARP curves for the Knee-Hip joint pair. Shaded vertical bands represent areas with a significant difference between curves and are coloured with the associated Cohen’s d effect size

Young children showed no significant differences compared to adolescents on the even surface but exhibited lower knee-hip MARP on the moderately uneven surface during initial contact and loading (0-5%, p<0.001, d=1.691 – large effect; 13-25%, p=0.001, d=1.466 – large effect). This more in-phase coupling was maintained on the highly uneven surface and extended to cover the initial swing (0-40%, p<0.001, d=1.902 – large effect; 61-84%, p<0.001, d=1.922 – large effect). Young children showed a greater in-phase pattern than children during initial swing on the highly uneven surface (53-80%, p<0.001, d=1.788 – large effect), with no differences observed on even and moderately uneven surfaces.

Finally, children showed lower in-phase coordination around toe-off on the even (56-65%, p=0.001, d=1.267 – large effect) and moderately uneven (53-67%, p<0.001, d=1.456 – large effect) surfaces compared to adults.

##### 3.2.1.2. Deviation phase

All groups showed a greater coordination variability (i.e., higher DP) when walking on uneven surfaces, with increases predominantly observed during initial contact and terminal swing, as displayed in Figure 5. For example, when comparing the highly uneven to the even surface, adults exhibited greater variability through the entire gait cycle (0-100%, p<0.001, d=2.487 – large effect), while in adolescents, children and young children, differences between these two surfaces were observed during initial contact (Adolescents: 0-42%, p<0.001, d=1.545 – large effect; Children: 0-15%, p<0.001, d=1.664 – large effect; Young children: 0-15%, p<0.001, d=1.893 – large effect), midstance (Adolescents: 0-42%, p<0.001, d=1.545 – large effect; Children: 23-50%, p<0.001, d=1.137 – large effect) and swing (Adolescents: 60-100%, p<0.001, d=2.106 – large effect; Children: 68-100%, p<0.001, d=1.314 – large effect; Young children: 68-100%, p<0.001, d=1.666 – large effect).

**Figure 5:**
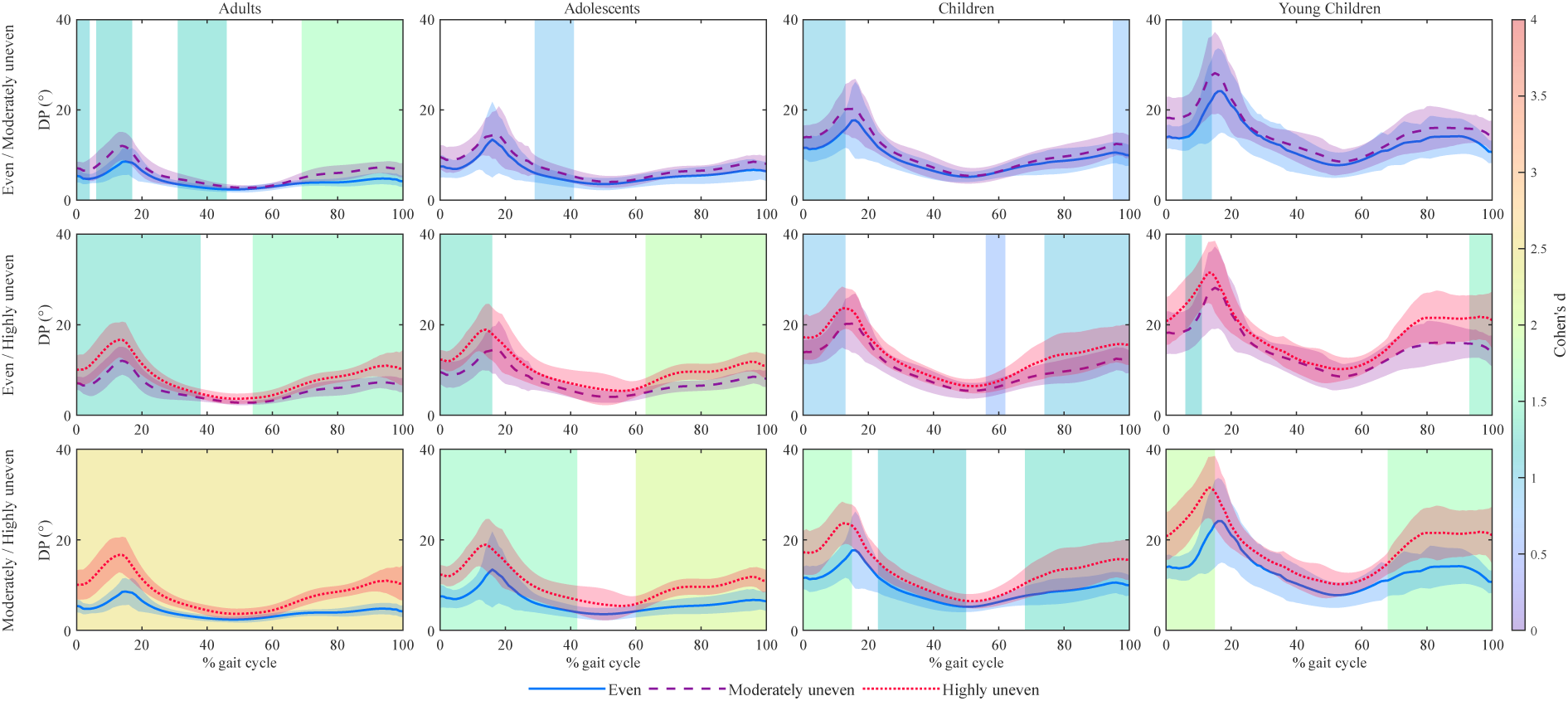
Surface comparisons of DP curves for the Knee-Hip joint pair. Shaded vertical bands represent areas with a significant difference between curves and are coloured with the associated Cohen’s d effect size

When comparing age groups (Figure 6), DP was higher in younger participants. Specifically, children and young children showed greater variability than adults over the entire gait cycle across all three surfaces (Even: Children, p<0.001, d=2.700 – large effect; Young children, p<0.001, d=3.158 – large effect; Moderately uneven: Children, p<0.001, d=2.207 – large effect; Young children, p<0.001, d=3.362 – large effect; Highly uneven: Children, p<0.001, d=1.907 – large effect; Young children, p<0.001, d=3.484 – large effect). Adolescents showed lower DP than children (Even: 0-13%, p<0.001, d=1.691 - large effect, 28-100%, p<0.001, d=1.427 - large effect; Moderately uneven: 0-14%, p<0.001, d=1.759 - large effect, 63-100%, p<0.001, d=1.513 - large effect; Highly uneven: 5-11%, p=0.002, d=1.298 - large effect) and young children (Even: 0-15%, p<0.001, d=2.333 - large effect; 17-100%, p<0.001, d=2.119 - large effect; Moderately uneven: 0-100%, p<0.001, d=2.479 - large effect; Highly uneven: 0-100%, p<0.001, d=2.357 - large effect) across the three surfaces, and significant differences were also observed when comparing adults to adolescents (Even: 1-6%, p<0.001, d=1.339 - large effect, 24-48%, p<0.001, d=1.438 - large effect; Moderately uneven: 22-64%, p<0.001, d=1.482 - large effect; Highly uneven: 26-40%, p<0.001, d=1.313 - large effect) and children to young children (Even: 22-36%, p<0.001, d=1.356 - large effect, 44-50%, p=0.002, d=1.303 – large effect, 74-92%, p<0.001, d=1.390 – large effect; Moderately uneven: 23-95%, p<0.001, d=1.567 - large effect; Highly uneven: 19-98%, p<0.001, d=1.623 – large effect), with the younger group showing greater variability in both cases (Figure 6).

**Figure 6:**
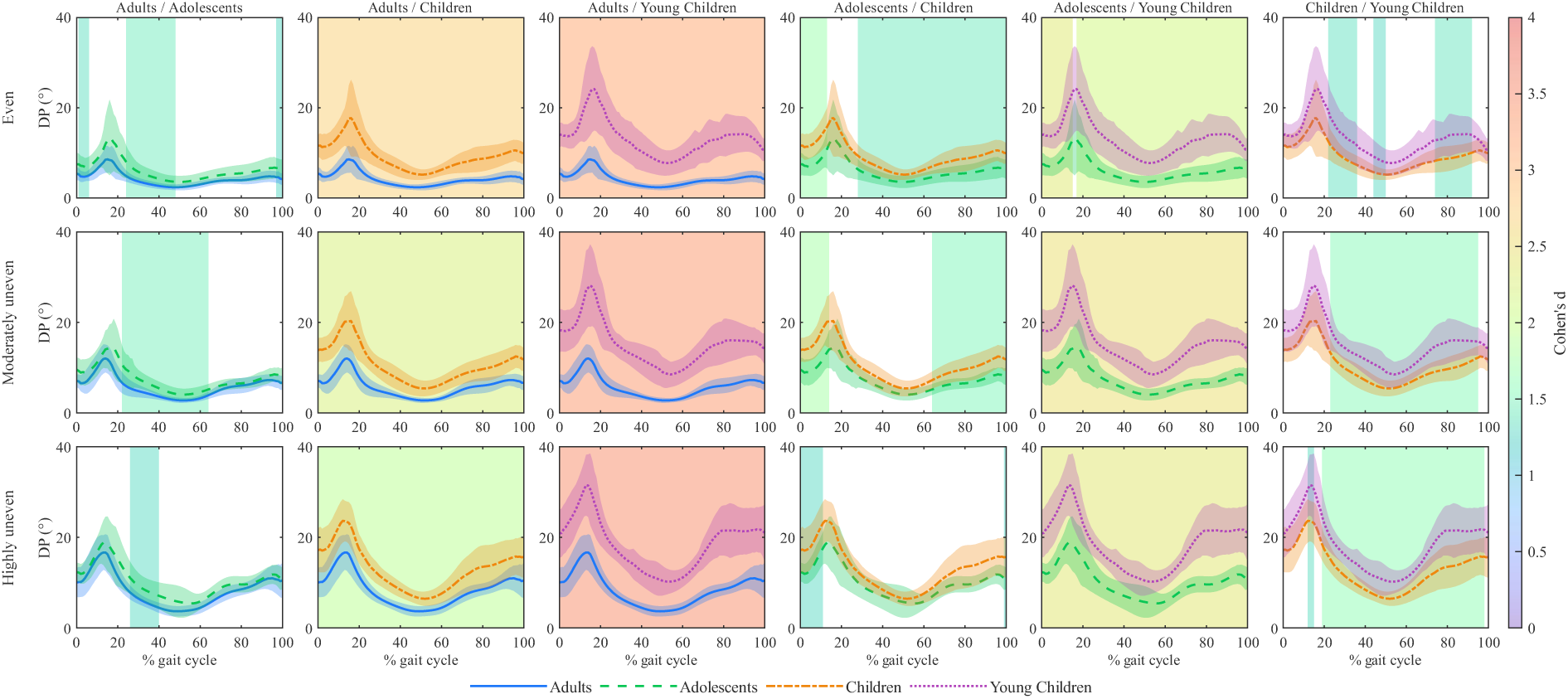
Group comparisons of DP curves for the Knee-Hip joint pair. Shaded vertical bands represent areas with a significant difference between curves and are coloured with the associated Cohen’s d effect size

#### 3.2.2. Ankle-Knee coordination

##### 3.2.2.1. Mean absolute relative phase

When comparing the highly uneven and the even surfaces, adults, adolescents and children showed a more in-phase ankle-knee coupling during swing when walking on the highly uneven surface (Adults: 56-99%, p<0.001, d=1.280 – large effect; Adolescents: 60-85%, p<0.001, d=1.099 – large effect; Children: 56-85%, p<0.001, d=1.536 – large effect; 86-100%, p<0.001, d=0.944 – large effect). For young children, a more in-phase coordination was also observed (Figure 7) during stance, initial and terminal swing (24-76%, p<0.001, d=1.483 – large effect, 92-100%, p<0.001, d=1.116 – large effect).

**Figure 7:**
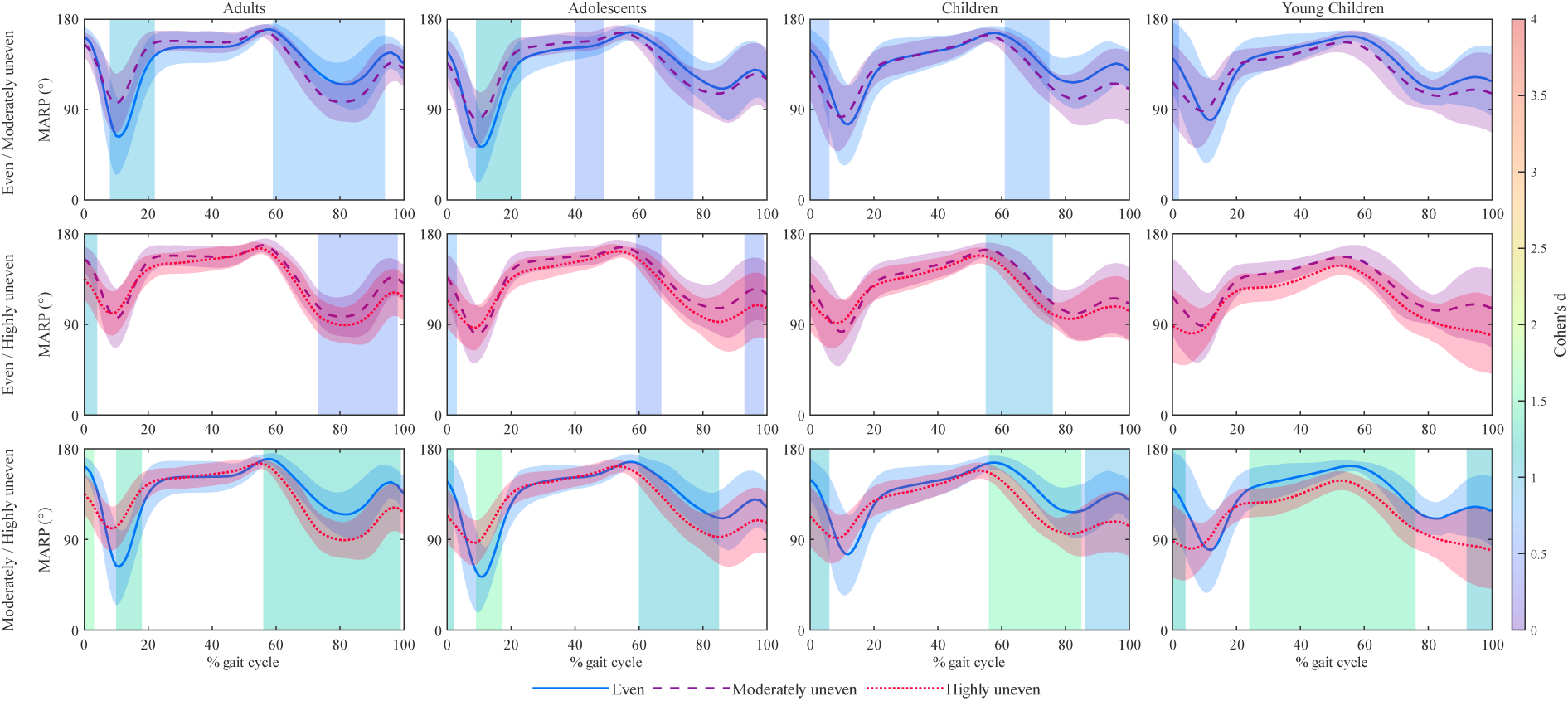
Surface comparisons of MARP curves for the Ankle-Knee joint pair. Shaded vertical bands represent areas with a significant difference between curves and are coloured with the associated Cohen’s d effect size

When comparing the moderately uneven and the even surfaces, a more in-phase coordination was observed during swing in adults (59-94%, p<0.001, d=0.827 – large effect), and early swing in adolescents (65-77%, p<0.001, d=0.692 – medium effect) and children (61-75%, p<0.001, d=0.713 – medium effect) when walking on the moderately uneven surface.

In a similar fashion, a more in-phase ankle-knee coupling was observed on the highly uneven surface compared to the moderately uneven surface during swing in adults (73-98%, p<0.001, d=0.508 – medium effect), adolescents (59-67%, p<0.001, d=0.507 – medium effect; 93-99%, p=0.003, d=0.507 – medium effect) and children (55-76%, p<0.001, d=0.938 – large effect).

In comparison to the even surface, adults and adolescents showed a more out-of-phase ankle-knee coordination during the loading phase when walking on uneven surfaces (Moderately uneven: Adults, 8-22%, p<0.001, d=1.022 – large effect; Adolescents, 9-23%, p<0.001, d=1.129 – large effect; Highly uneven: Adults, 10-18%, p<0.001, d=1.389 – large effect; Adolescents, 9-17%, p<0.001, d=1.560 – large effect). For adolescents, this more out-of-phase coupling also occurred during mid stance when comparing the moderately uneven with the even surface (40-49%, p=0.003, d=0.537 – medium effect).

Regarding age group differences (Figure 8), children and young children exhibited a more in-phase ankle-knee coordination than adults during stance only when walking on uneven surfaces (Moderately uneven: Children, 18-32%, p<0.001, d=1.419 – large effect; Young children, 20-37%, p<0.001, d=1.575 – large effect; Highly uneven: Children, 20-29%, p=0.001, d=1.268 – large effect; Young children, 18-61%, p<0.001, d=1.807 – large effect). Young children also showed a more in-phase coordination than adolescents (28-43%, p<0.001, d=1.534 – large effect, 49-60%, p<0.001, d=1.674 – large effect) during stance when walking on the highly uneven surface.

**Figure 8:**
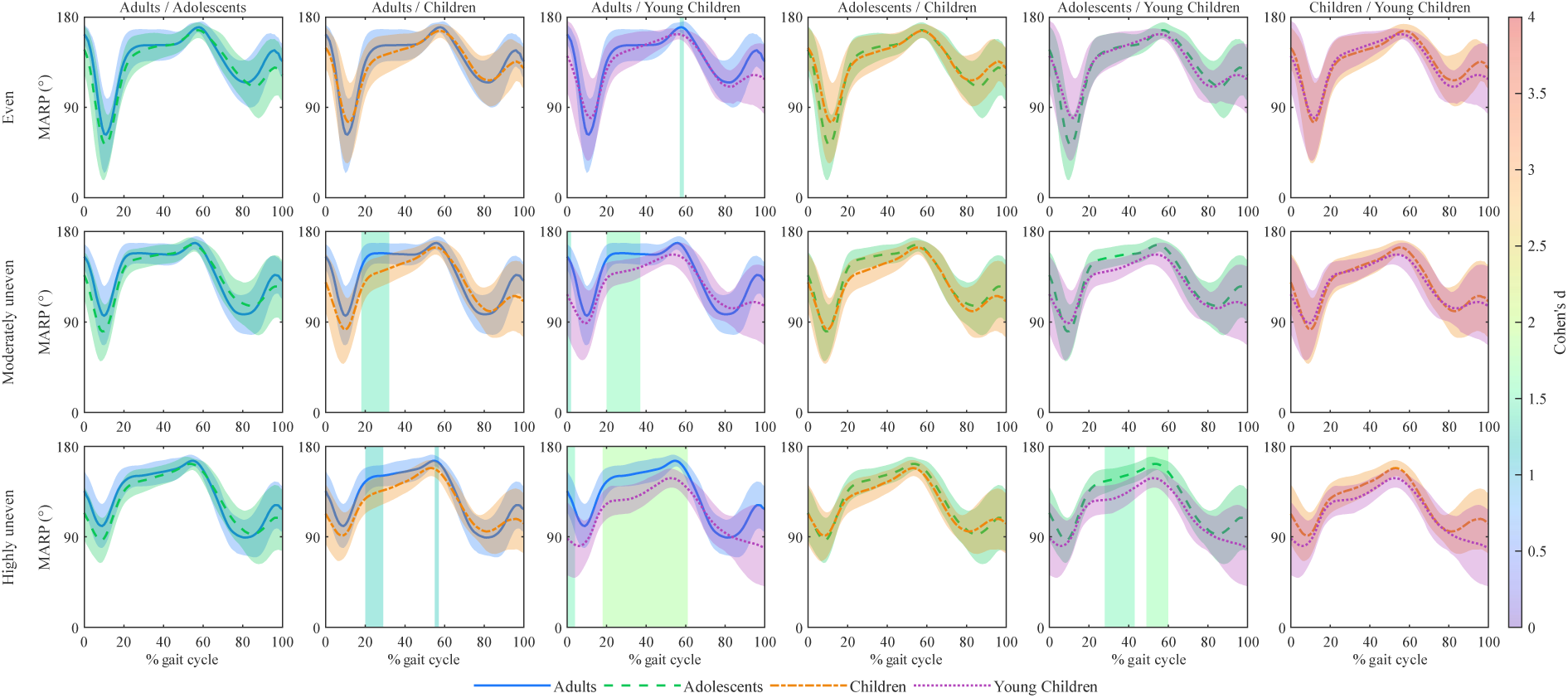
Group comparisons of MARP curves for the Ankle-Knee joint pair. Shaded vertical bands represent areas with a significant difference between curves and are coloured with the associated Cohen’s d effect size

##### 3.2.2.2. Deviation phase

A greater ankle-knee coordination variability was observed in all groups when walking on uneven surfaces (Figure 9). Notably, a higher DP was observed in all groups throughout nearly the entire gait cycle (Adults: 0-100%, p<0.001, d=2.203 – large effect, Adolescents: 0-7%, p=0.002, d=1.394 – large effect, 8-80%, p<0.001, d=2.196 – large effect, 91-100%, p=0.001, d=1.511 – large effect; Children: 9-100%, p<0.001, d=2.017 – large effect; Young children: 11-17%, p=0.003, d=1.655 – large effect, 23-83%, p<0.001, d=2.076 – large effect) when comparing the highly uneven surface to the even one.

**Figure 9:**
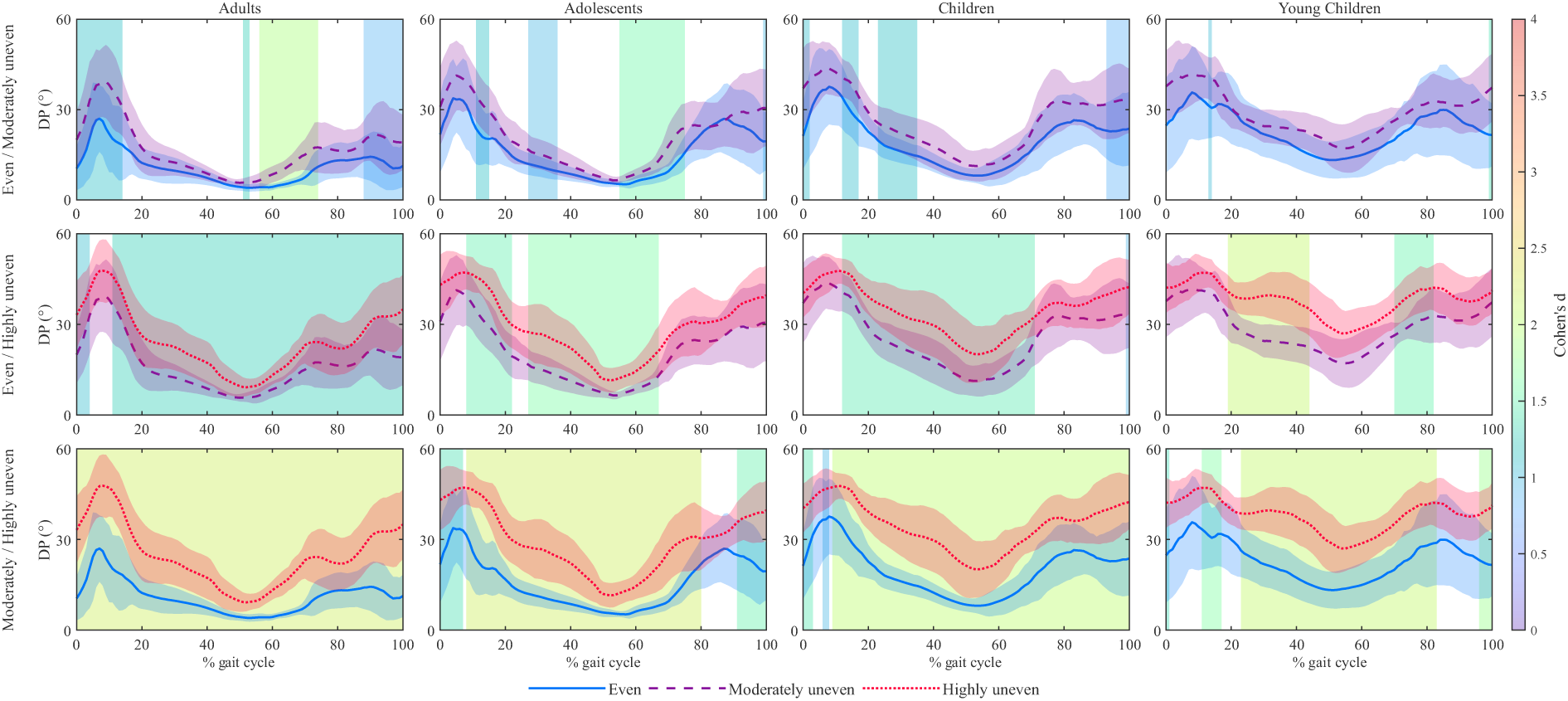
Group comparisons of DP curves for the Ankle-Knee joint pair. Shaded vertical bands represent areas with a significant difference between curves and are coloured with the associated Cohen’s d effect size

When comparing age groups, DP was greater in younger participants across the three surfaces, with statistically significant differences appearing mostly when comparing adults with children (Even: 15-90%, p<0.001, d=1.711 – large effect; Moderately uneven: 15-57%, p<0.001, d=1.652 - large effect, 73-90%, p<0.001, d=1.753; Highly uneven: 15-72%, p<0.001, d=1.537 - large effect, 72-87%, p<0.001, d=1.559 - large effect) and young children (Even: 15-88%, p<0.001, d=2.030 - large effect; Moderately uneven: 16-89%, p<0.001, d=2.367 - large effect; Highly uneven: 17-88%, p<0.001, d=2.454 - large effect), as well as adolescents with young children (Even: 18-72%, p<0.001, d=1.776 – large effect; Moderately uneven: 23-70%, p<0.001, d=2.067 - large effect; Highly uneven: 23-68%, p<0.001, d=1.962 - large effect) (Figure 10).

**Figure 10:**
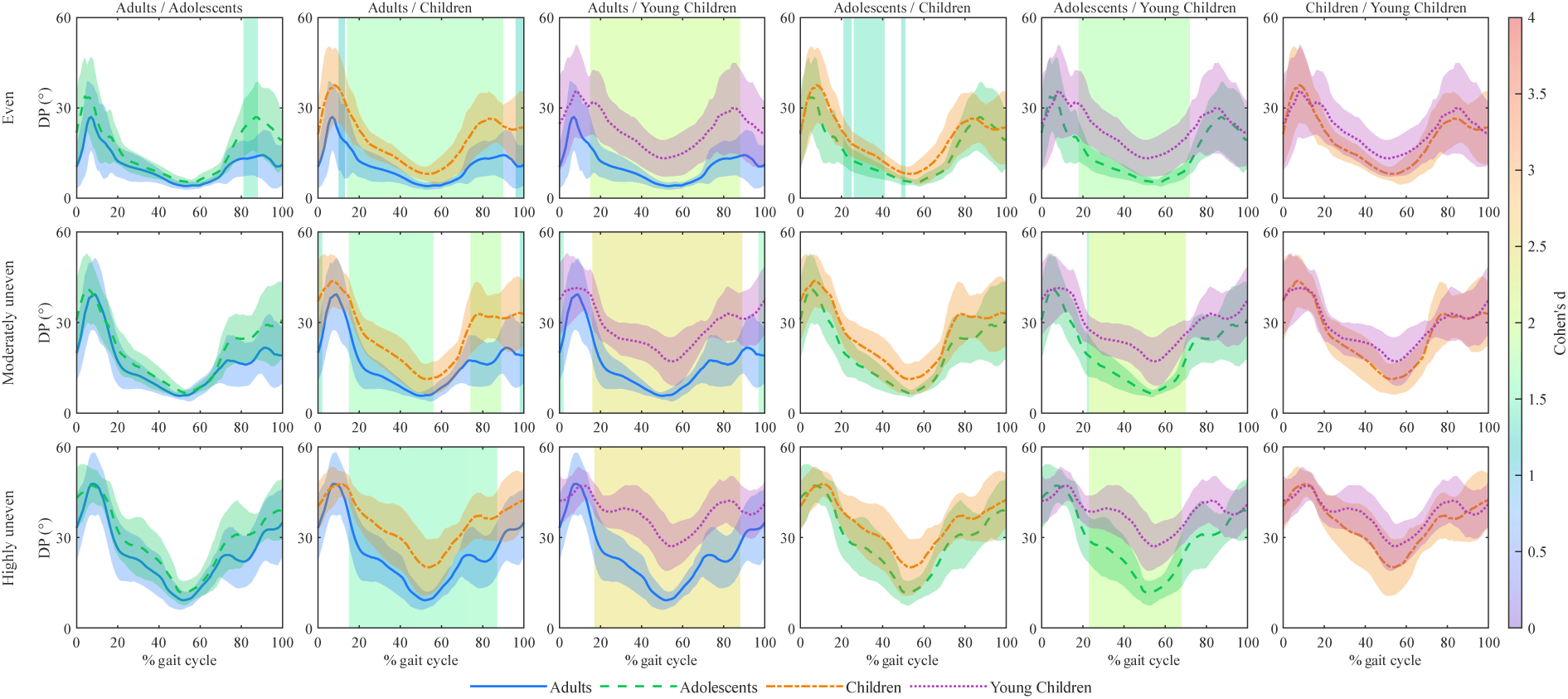
Group comparisons of DP curves for the Ankle-Knee joint pair. Shaded vertical bands represent areas with a significant difference between curves and are coloured with the associated Cohen’s d effect size

## 4. Discussion

This study aimed to assess the adaptations in intersegmental and inter-joint coordination when walking on uneven surfaces, to quantify inter-joint coordination variability, and to investigate the differences in adaptation between age groups on uneven surfaces. Overall, the findings support our hypotheses, showing that uneven surfaces affect coordination patterns, leading to more in-phase inter-joint coordination, less dense kinectomes and increased variability. These variables also differ across age groups, with younger participants exhibiting more in-phase inter-joint coordination and older participants showing denser kinectomes.

Four main results were observed. First, the organization of intersegmental coordination was overall preserved across age groups and surfaces, while kinectome density increased with age and decreased with surface irregularity. Second, for the inter-joint approach, results showed a more in-phase knee-hip and ankle-knee coordination on both the moderately and highly uneven surfaces for all age groups, in comparison to the even surface. Third, younger participants exhibited a more in-phase inter-joint coordination than older participants, and this tendency was emphasized on uneven surfaces. Fourth, coordination variability, as measured by DP, was also higher in younger groups regardless of the surface.

### 4.1. Intersegmental coordination strategies

The identification of stable marker communities across age groups and surfaces (Figure 1) suggests that the fundamental organisation of intersegmental coordination emerges early and remains robust throughout development. In particular, the clustering into a head-trunk unit and two contralateral upper and lower limb communities reflects fundamental patterns of human locomotion, as head-trunk stability has been shown to emerge between 6 and 32 weeks of independent walking (Ledebt & Bril, 2000), while reciprocal arm swing is typically present by 18 months of age (Sutherland et al., 1980). Although previous work has reported age-related changes in intersegmental coordination during gait using measures such as MARP between ipsilateral and contralateral upper and lower limbs (Meyns et al., 2020), the present study extends these observations at the whole-body level using kinectome analysis. The observed increase in kinectome density with age may therefore reflect a progressive strengthening and refinement of intersegmental coordination throughout maturation, highlighting the use of kinectome density as a promising indicator of locomotor development. Finally, the decrease in network density observed across all age groups with increasing irregularities suggests a disruption in these coordinated interactions in response to environmental constraints. Rather than reflecting a simple decline in motor control, this decrease may indicate a functional reorganization of coordination in which segmental interactions become more flexible and less constrained to accommodate external perturbations (i.e., irregular surfaces) (Kent et al., 2019).

### 4.2. Inter-joint coordination on uneven surfaces

For CRP analysis, the more in-phase knee-hip and ankle-knee coordination observed in all age groups during stance (knee-hip only) and swing (knee-hip and ankle-knee) on uneven surfaces (Figures 3 and 7) align with earlier research. For instance, one study has shown more in-phase knee-hip and ankle-knee coordination in children with cerebral palsy and typically developing peers during the swing phase, when walking on an uneven surface (Dussault-Picard et al., 2023). Another study has reported more in-phase knee-hip coordination in all phases of the gait cycle and more in-phase ankle-knee coupling during initial contact, loading and swing when young and older adults walked on an irregular brick surface compared to a flat walkway (Ippersiel et al., 2021). In line with the hypotheses of these two studies, we suggest that the more in-phase joint coordination observed on uneven surfaces results from a more cautious motor strategy adopted to reduce the risk of balance loss. When comparing the uneven and even surfaces, more in-phase ankle-knee coordination was mainly observed during the swing phase, which suggests that inter-joint coupling strengthened in preparation for ground contact and to ensure sufficient toe-clearance over the irregularities. This interpretation is consistent with previous work reporting higher minimum toe clearance on uneven terrain (Schulz, 2011) and increased pre-activation in lower leg muscles in late swing (Müller et al., 2010), reflecting an anticipatory control strategy that may contributes to increased ankle stiffness at foot contact on uneven ground. More in-phase knee-hip patterns occurred during both stance and swing, which conveys the idea that the more rigid coordination between these joints reflects, in addition to preparatory adjustments for foot contact, a strategy aimed at maintaining a stable posture on uneven ground. This is in line with reports of increased co-contractions during stance on uneven surfaces (Voloshina et al., 2013) and evidence of increased quasi-stiffness of the knee joint on destabilizing ground (Foster et al., 2020). Concerning the more out-of-phase ankle-knee coordination observed during loading on the uneven surfaces, we hypothesised that it resulted from modulations of the coordination pattern caused by the foot position on surface irregularities. In this context, the ankle was directly influenced by local surface features while the knee tended to preserve its habitual loading pattern. Joint kinematic signals and variability, which support this hypothesis, are available in the Supplementary Material.

### 4.3. Age-related changes in inter-joint coordination

Differences in motor control between age groups are highlighted by the more in-phase coordination observed in younger participants (Figures 4 and 8). This pattern suggests a more rigid motor control, often interpreted as a freezing of degrees of freedom to simplify the motor task (Ko et al., 2003; Vereijken et al., 1992). However, young children also showed greater coordination variability (i.e., higher DP) than adults and adolescents for both joint pairs (Figures 6 and 10). Together, these two results (lower MARP and higher DP) suggest a less mature and inherently less stable gait pattern, indicating that although motor control is simplified, young children are not yet able to maintain this coupling consistently over time. This interpretation is in line with the concept of block-like coordination in early development, where movement is produced through rigidly linked segments, reducing motor complexity (Assaiante, 1998; Dominici et al., 2011). Also, it has been shown that gait variability is higher in young children and decreases throughout development (Hausdorff et al., 1999; Sutherland et al., 1980). Regarding the more out-of-phase knee-hip coordination observed around toe-off when comparing children to adults, we hypothesized that it resulted from slight temporal shifts in angular minima and maxima throughout the gait cycle between the two groups (Supplementary material: Figures S3 and S4). The immaturity and instability of the gait pattern is further highlighted by the fact that coordination differences between age groups were emphasised by surface irregularities. Young participants exhibited more in-phase ankle-knee coordination than older participants on uneven surfaces during the stance phase (Figure 8), suggesting that they rely more on distal coupling to maintain stability on uneven ground. Regarding proximal coordination, young children also exhibited more in-phase knee-hip coupling than adults and adolescents during stance on both uneven surfaces, supporting a more globally coupled organization of lower-limb coordination under increased balance demands. Thus, the more in-phase coordination observed in younger participants on uneven surfaces may indicate a more cautious motor strategy relying on increased joint coupling to enhance stability, likely reflecting ongoing maturation of neuromotor control. While locomotion is mainly governed by spinal circuits which generate cyclical and rhythmic movement patterns, the maturation of motor control is accompanied by a greater implication of the corticospinal tract, the pathway responsible for voluntary movements (Salehpour et al., 2026). This maturation leads to a more effective modulation of the spinal patterns and thus a greater adaptivity to challenging environments (Salehpour et al., 2026), which might explain why differences in gait coordination across age groups were more pronounced on the uneven surfaces.

### 4.4. Deviation phase

In this context, the higher coordinative variability observed in young children should not be viewed solely as a lack of control, but rather as a necessary part of neuromotor development (Hadders-Algra, 2010). Indeed, a high level of variability is typical during the learning phase (Dusing & Harbourne, 2010; Stergiou & Decker, 2011), providing the flexibility necessary to explore different coordination strategies. The increase in variability on uneven surfaces was observed across all age groups (Figures 5 and 9), which highlights that challenging environments can affect gait consistency regardless of motor maturity. However, the distinction lies in the nature of this response, as it likely reflects a progressive shift from predominantly exploratory variability in younger participants (Hadders-Algra, 2010) to increasingly functional adaptability with neuromotor maturation (Stergiou & Decker, 2011).

### 4.5. Limitations

The results presented must be interpreted considering several limitations. First, participants were classified into four age groups, which may not adequately capture the continuous nature of maturation. Indeed, the children and adolescent groups span relatively broad age ranges (6-11 and 12-17 years old respectively) involving substantial neuromotor and growth-related changes (Bisi & Stagni, 2016; Largo & Rousson, 2003) that may not be fully reflected by categorical grouping. Second, the CRP was only computed from sagittal joint angles, and kinectomes were built from marker accelerations along the anteroposterior axis. Therefore, the results presented do not capture potential gait differences occurring along other axes or in other planes of movement. This choice was motivated by the fact that these dimensions involve smaller ranges of motion during gait and are therefore less likely to contribute substantially to the characterization of coordination. Moreover, joint angles in the frontal and transverse planes are more prone to measurement error (Fonseca et al., 2022), which may introduce bias in CRP values. Third, walking was performed at a self-selected speed, which may have influenced coordination outcomes (Fonseca et al., 2022). Fourth, the number of trials differed between groups, with young children completing fewer trials than older participants. Although this was done to limit fatigue, it may have affected the MARP and DP estimates (Dussault-Picard et al., 2026). Finally, CRP is inherently sensitive to signal processing choices (Ippersiel et al., 2019; Lamb & Stöckl, 2014) and assessor-dependent factors. Previous research has shown that intra-therapist reliability of CRP measurements is subject to variability (Dussault-Picard et al., 2026), which may have influenced the results of group comparisons.

## 5. Conclusion

The results of the present study showed that uneven surfaces induce changes in gait coordination, leading to more in-phase inter-joint coupling, reduced kinectome density and greater inter-joint coordination variability. The more in-phase inter-joint coupling observed on uneven surfaces likely reflects a cautious motor strategy aimed at maintaining postural stability. Regarding age comparisons, stable marker communities across age groups suggest an early establishment of the architecture of whole-body segmental coordination, while the age-related increase in kinectome density likely reflects stronger interactions between segments with maturation. Younger participants showed more in-phase coordination and greater variability, reflecting less mature motor control. Finally, age-related differences in inter-joint coordination were emphasized on uneven surfaces, likely reflecting maturation-related improvements in supraspinal modulation of spinal locomotor patterns and, consequently, greater adaptability to environmental perturbations. From a clinical perspective, these findings may contribute to a better understanding of gait stability and motor control strategies in challenging environments and could inform the assessment and rehabilitation of individuals with impaired locomotor adaptability.

## Author contributions

Conceptualization: AC, CDP, YC; Investigation: AC, SDF; Data curation: AC, SDF; Formal analysis: AC; Software: AC; Funding acquisition: YC; Supervision: YC; Visualization: AC; Writing – original draft: AC, CDP; Writing review & editing: AC, CDP, SDF, YC.

## Data availability statement

The data used in this study can be obtained from the corresponding author upon reasonable request and with permission of the ethics committee of the Centre de Recherche Azrieli du CHU Sainte-Justine.

## Declaration of generative AI use

Generative AI tools were used to assist with language editing and manuscript rephrasing.

## Abbreviations

CRP: Continuous Relative Phase
MARP: Mean Absolute Relative Phase
DP: Deviation phase
SPM: Statistical Parametric Mapping

## References

Assaiante, C. (1998). Development of Locomotor Balance Control in Healthy Children. Neuroscience & Biobehavioral Reviews, 22(4), 527-532. 10.1016/S0149-7634(97)00040-7

Bisi, M. C., & Stagni, R. (2016). Development of gait motor control: What happens after a sudden increase in height during adolescence? BioMedical Engineering OnLine, 15, 47. 10.1186/s12938-016-0159-0

Blondel, V. D., Guillaume, J.-L., Lambiotte, R., & Lefebvre, E. (2008). Fast unfolding of communities in large networks. Journal of Statistical Mechanics: Theory and Experiment, 2008(10), P10008. 10.1088/1742-5468/2008/10/P10008

Brognara, L., Arceri, A., Zironi, M., Traina, F., Faldini, C., & Mazzotti, A. (2025). Gait Spatio-Temporal Parameters Vary Significantly Between Indoor, Outdoor and Different Surfaces. Sensors, 25(5), 1314. 10.3390/s25051314

Burgess-Limerick, R., Abernethy, B., & Neal, R. J. (1993). Relative phase quantifies interjoint coordination. Journal of Biomechanics, 26(1), 91-94. 10.1016/0021-9290(93)90617-N

Carollo, J. J., Worster, K., Pan, Z., Ma, J., Chang, F., & Valvano, J. (2018). Relative phase measures of intersegmental coordination describe motor control impairments in children with cerebral palsy who exhibit stiff-knee gait. Clinical Biomechanics, 59, 40-46. 10.1016/j.clinbiomech.2018.07.015

Celestino, M. L., van Emmerik, R., Barela, J. A., Gama, G. L., & Barela, A. M. F. (2019). Intralimb gait coordination of individuals with stroke using vector coding. Human Movement Science, 68, 102522. 10.1016/j.humov.2019.102522

Cohen, J. (avec Internet Archive). (1988). Statistical power analysis for the behavioral sciences. Hillsdale, N.J.: L. Erlbaum Associates. http://archive.org/details/statisticalpower0000cohe_j0l3

Dominici, N., Ivanenko, Y. P., Cappellini, G., d’Avella, A., Mondì, V., Cicchese, M., Fabiano, A., Silei, T., Di Paolo, A., Giannini, C., Poppele, R. E., & Lacquaniti, F. (2011). Locomotor Primitives in Newborn Babies and Their Development. Science, 334(6058), 997-999. 10.1126/science.1210617

Dunn, O. J. (1961). Multiple Comparisons among Means. Journal of the American Statistical Association, 56(293), 52-64. 10.1080/01621459.1961.10482090

Dusing, S. C., & Harbourne, R. T. (2010). Variability in Postural Control During Infancy: Implications for Development, Assessment, and Intervention. Physical Therapy, 90(12), 1838-1849. 10.2522/ptj.2010033

Dussault-Picard, C., Cherni, Y., & Dixon, P. C. (2025). Spatiotemporal characteristics of gait when walking on an uneven surface in children with cerebral palsy. Scientific Reports, 15(1), 4912. 10.1038/s41598-025-89280-x

Dussault-Picard, C., Cherni, Y., Ferron, A., Robert, M. T., & Dixon, P. C. (2023). The effect of uneven surfaces on inter-joint coordination during walking in children with cerebral palsy. Scientific Reports, 13(1), 21779. 10.1038/s41598-023-49196-w

Dussault-Picard, C., Cherni, Y., Fonseca, M., Carcreff, L., Leboeuf, F., & Armand, S. (2026). Intra-and inter-therapist reliability of lower-limb inter-joint coordination during gait in individuals with and without cerebral palsy. Gait & Posture, 127, 110148. 10.1016/j.gaitpost.2026.110148

Dussault-Picard, C., Ippersiel, P., Böhm, H., & Dixon, P. C. (2022). Lower-limb joint-coordination and coordination variability during gait in children with cerebral palsy. Clinical Biomechanics, 98, 105740. 10.1016/j.clinbiomech.2022.105740

Dussault-Picard, C., Mohammadyari, S. G., Arvisais, D., Robert, M. T., & Dixon, P. C. (2022). Gait adaptations of individuals with cerebral palsy on irregular surfaces: A scoping review. Gait & Posture, 96, 35-46. 10.1016/j.gaitpost.2022.05.011

Fonseca, M., Bergere, M., Candido, J., Leboeuf, F., Dumas, R., & Armand, S. (2022). The Conventional Gait Model’s sensitivity to lower-limb marker placement. Scientific Reports, 12(1), 14207. 10.1038/s41598-022-18546-5

Foster, A. J., Hudson, P. E., & Smith, N. (2020). Quasi-stiffness of the knee joint is influenced by walking on a destabilising terrain. The Knee, 27(6), 1889-1898. 10.1016/j.knee.2020.09.019

Froehle, A. W., Nahhas, R. W., Sherwood, R. J., & Duren, D. L. (2013). Age-related changes in spatiotemporal characteristics of gait accompany ongoing lower limb linear growth in late childhood and early adolescence. Gait & Posture, 38(1), 14-19. 10.1016/j.gaitpost.2012.10.005

Gentle, J., Barnett, A. L., & Wilmut, K. (2016). Adaptations to walking on an uneven terrain for individuals with and without Developmental Coordination Disorder. Human Movement Science, 49, 346-353. 10.1016/j.humov.2016.08.010

Goetschalckx, M., Moumdjian, L., Feys, P., & Rameckers, E. (2024). Interlimb coordination and spatiotemporal variability during walking and running in children with developmental coordination disorder and typically developing children. Human Movement Science, 96, 103252. 10.1016/j.humov.2024.103252

Gómez, S., Jensen, P., & Arenas, A. (2009). Analysis of community structure in networks of correlated data. Physical Review E, 80(1), 016114. 10.1103/PhysRevE.80.016114

Hadders-Algra, M. (2010). Variation and Variability: Key Words in Human Motor Development. Physical Therapy, 90(12), 1823-1837. 10.2522/ptj.20100006

Hamill, J., van Emmerik, R. E. A., Heiderscheit, B. C., & Li, L. (1999). A dynamical systems approach to lower extremity running injuries. Clinical Biomechanics, 14(5), 297-308. 10.1016/S0268-0033(98)90092-4

Hausdorff, J. M., Zemany, L., Peng, C.-K., & Goldberger, A. L. (1999). Maturation of gait dynamics: Stride-to-stride variability and its temporal organization in children. Journal of Applied Physiology, 86(3), 1040-1047. 10.1152/jappl.1999.86.3.1040

Heiderscheit, B. C., Hamill, J., & van Emmerik, R. E. A.. (2002). Variability of Stride Characteristics and Joint Coordination among Individuals with Unilateral Patellofemoral Pain. Journal of Applied Biomechanics, 18(2), 110-121. 10.1123/jab.18.2.110

Inns, T. B., Pina, I., Macgregor, L. J., Dudchenko, P. A., Crockett, R. A., & Hunter, A. M. (2025). Age-related gait adaptations: Analysis of temporal gait parameters and variability, and muscle activation across flat vs. uneven surfaces in young, middle-aged, and older adults. Frontiers in Aging, 6. 10.3389/fragi.2025.1573778

Ippersiel, P., Preuss, R., & Robbins, S. M. (2019). The Effects of Data Padding Techniques on Continuous Relative-Phase Analysis Using the Hilbert Transform. Journal of Applied Biomechanics, 35(4), 247-255. 10.1123/jab.2018-0396

Ippersiel, P., Robbins, S. M., & Dixon, P. C. (2021). Lower-limb coordination and variability during gait: The effects of age and walking surface. Gait & Posture, 85, 251-257. 10.1016/j.gaitpost.2021.02.009

Ippersiel, P., Shah, V., & Dixon, P. C. (2022). The impact of outdoor walking surfaces on lower-limb coordination and variability during gait in healthy adults. Gait & Posture, 91, 7-13. 10.1016/j.gaitpost.2021.09.176

Kelso, J. A. S. (1995). Dynamic patterns: The self-organization of brain and behavior (p. xvii, 334). The MIT Press.

Kent, J. A., Sommerfeld, J. H., Mukherjee, M., Takahashi, K. Z., & Stergiou, N. (2019). Locomotor patterns change over time during walking on an uneven surface. Journal of Experimental Biology, 222(14), jeb202093. 10.1242/jeb.202093

Ko, Y.-G., Challis, J. H., & Newell, K. M. (2003). Learning to coordinate redundant degrees of freedom in a dynamic balance task. Human Movement Science, 22(1), 47-66. 10.1016/S0167-9457(02)00177-X

Lally, M., & Valentine-French, S. (2017). *Lifespan Development: A Psychological Perspective* (2nd éd.). Open textbook.

Lamb, P. F., & Stöckl, M. (2014). On the use of continuous relative phase: Review of current approaches and outline for a new standard. Clinical Biomechanics, 29(5), 484-493. 10.1016/j.clinbiomech.2014.03.008

Largo, R. H., & Rousson, V. (2003). Neuromotor development from kindergarten age to adolescence: Developmental course and variability. Swiss Medical Weekly, 133(1314), 193-199. 10.4414/smw.2003.09883

Ledebt, A., & Bril, B. (2000). Acquisition of upper body stability during walking in toddlers. Developmental Psychobiology, 36(4), 311-324. 10.1002/(SICI)1098-2302(200005)36:4%3C311::AID-DEV6%3E3.0.CO;2-V

Meyns, P., Van de Walle, P., Desloovere, K., Janssens, S., Van Sever, S., & Hallemans, A. (2020). Age-related differences in interlimb coordination during typical gait: An observational study. Gait & Posture, 81, 109-115. 10.1016/j.gaitpost.2020.07.013

Müller, R., Grimmer, S., & Blickhan, R. (2010). Running on uneven ground: Leg adjustments by muscle pre-activation control. Human Movement Science, 29(2), 299-310. 10.1016/j.humov.2010.01.003

Needham, R., Naemi, R., & Chockalingam, N. (2014). Quantifying lumbar–pelvis coordination during gait using a modified vector coding technique. Journal of Biomechanics, 47(5), 1020-1026. 10.1016/j.jbiomech.2013.12.032

Pataky, T. C. (2010). Generalized *n*-dimensional biomechanical field analysis using statistical parametric mapping. Journal of Biomechanics, 43(10), 1976-1982. 10.1016/j.jbiomech.2010.03.008

Pataky, T. C., Vanrenterghem, J., & Robinson, M. A. (2015). Zero-vs. One-dimensional, parametric vs. Non-parametric, and confidence interval vs. Hypothesis testing procedures in one-dimensional biomechanical trajectory analysis. Journal of Biomechanics, 48(7), 1277-1285. 10.1016/j.jbiomech.2015.02.051

Rubinov, M., Kötter, R., Hagmann, P., & Sporns, O. (2009). Brain connectivity toolbox: A collection of complex network measurements and brain connectivity datasets. NeuroImage, Organization for Human Brain Mapping 2009 Annual Meeting, 47, S169. 10.1016/S1053-8119(09)71822-1

Salehpour, S., Mathieu, S., Sylos-Labini, F., Ivanenko, Y., Lacquaniti, F., & Dewolf, A. H. (2026). A neuromechanical framework for the development of interlimb coordination. Neuroscience & Biobehavioral Reviews, 186, 106671. 10.1016/j.neubiorev.2026.106671

Schulz, B. W. (2011). Minimum toe clearance adaptations to floor surface irregularity and gait speed. Journal of Biomechanics, 44(7), 1277-1284. 10.1016/j.jbiomech.2011.02.010

Stergiou, N., & Decker, L. M. (2011). Human movement variability, nonlinear dynamics, and pathology: Is there a connection? Human Movement Science, EWOMS 2009: The European Workshop on Movement Science, 30(5), 869-888. 10.1016/j.humov.2011.06.002

Stergiou, N., Jensen, J. L., Bates, B. T., Scholten, S. D., & Tzetzis, G. (2001). A dynamical systems investigation of lower extremity coordination during running over obstacles. Clinical Biomechanics, 16(3), 213-221. 10.1016/S0268-0033(00)00090-5

Sutherland, D. H., Olshen, R., Cooper, L., & Woo, S. L. (1980). The development of mature gait. JBJS, 62(3), 336.

Troisi Lopez, E., Sorrentino, P., Liparoti, M., Minino, R., Polverino, A., Romano, A., Carotenuto, A., Amico, E., & Sorrentino, G. (2022). The kinectome: A comprehensive kinematic map of human motion in health and disease. Annals of the New York Academy of Sciences, 1516(1), 247-261. 10.1111/nyas.14860

van Emmerik, R. E. A., Miller, R. H., & Hamill, J. (2014). Dynamical Systems Analysis of Coordination. In Research Methods in Biomechanics (2e éd., p. 291-315). Human Kinetics. http://www.humankineticslibrary.com/hklp/encyclopedia-chapter

Vereijken, B., van Emmerik, R. E. A., Whiting, H. T. A., & Newell, K. M. (1992). Free(z)ing Degrees of Freedom in Skill Acquisition. Journal of Motor Behavior, 24(1), 133-142. 10.1080/00222895.1992.9941608

Voloshina, A. S., Kuo, A. D., Daley, M. A., & Ferris, D. P. (2013). Biomechanics and energetics of walking on uneven terrain. Journal of Experimental Biology, 216(21), 3963-3970. 10.1242/jeb.081711

