## Supplementary Material for "Impact of age and surface irregularities on intersegmental and inter-joint coordination during gait"

**Details of CRP calculations**

Joint kinematic signals from all gait trials were low-pass filtered (bidirectional Butterworth, 4th order, 6-Hz cut-off), segmented into cycles (consecutive heel strikes from the same foot), and padded with spline extrapolation over 10 points on both ends to avoid edge effects. Signals were then amplitude-centered around zero following equation (s1):

$$x_c(t) = x(t) - \min\big(x(t)\big) - \frac{\max\big(x(t)\big) - \min\big(x(t)\big)}{2}\ . \qquad (s1)$$

Phase angles were calculated using the Hilbert transform method. Specifically, centered signals $x_c(t)$ were transformed into analytic signals ζ(t) as in equation (s2):

$$\zeta(t) = x_c(t) + iH(t), \qquad (s2)$$

where the Hilbert transform H(t) is the imaginary part of ζ(t). The phase angle was then defined as the inverse tangent of the ratio between the imaginary and real parts of ζ(t) (Equation (s3)).

$$\phi(t) = \arctan\left(\frac{H(t)}{x_c(t)}\right) \qquad (s3)$$

Padding points were subsequently removed, and CRP was calculated for each cycle as the absolute difference between the phase angles of corresponding joint pairs (Equation (s4)).

$$CRP(t_i) = \phi_1(t) - \ \phi_2(t) \qquad (s4)$$

Finally, to address potential discontinuities across the gait cycle, CRP values above 180 were transformed by subtracting them from 360, thus mapping all values to a 0–180° range.

**Supplementary figures**

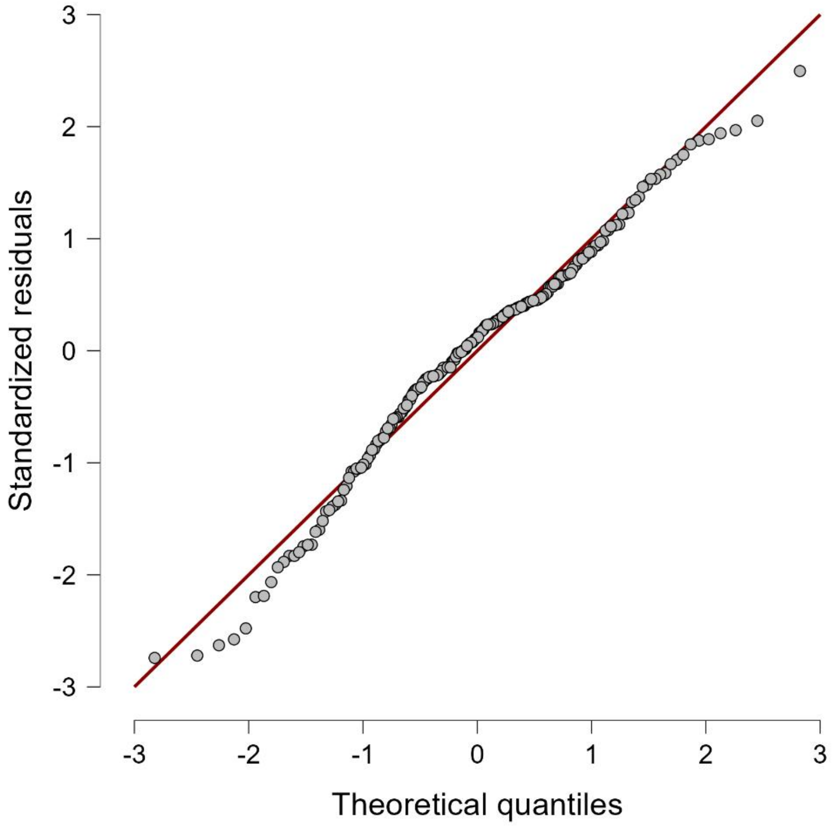


*Figure S1: Q-Q Plot of density residuals*

**Knee-Hip MARP**

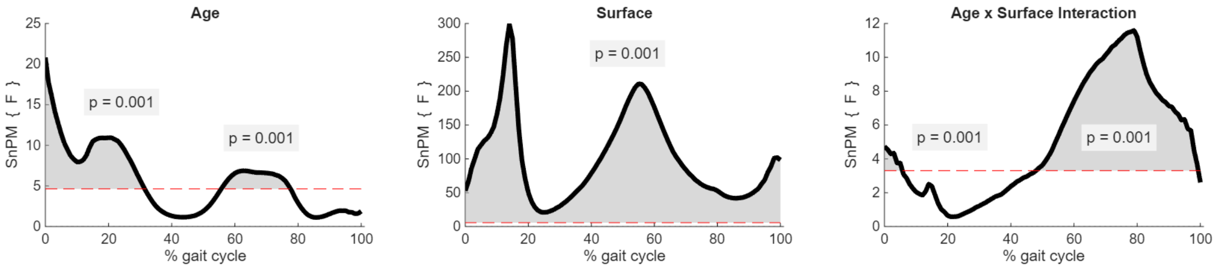


**Knee-Hip DP**

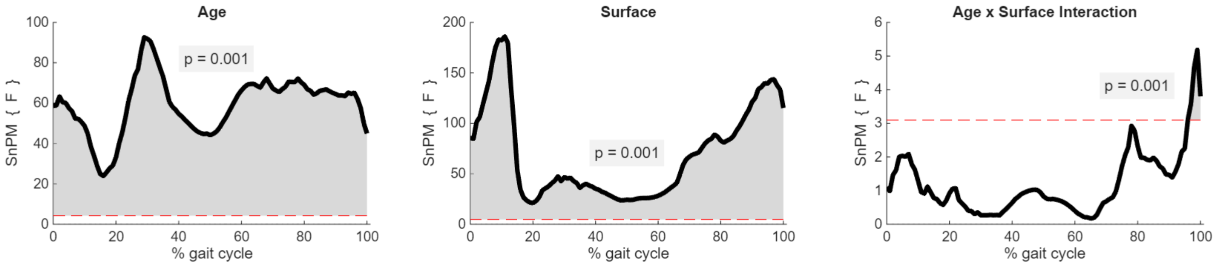


**Ankle-Knee MARP**

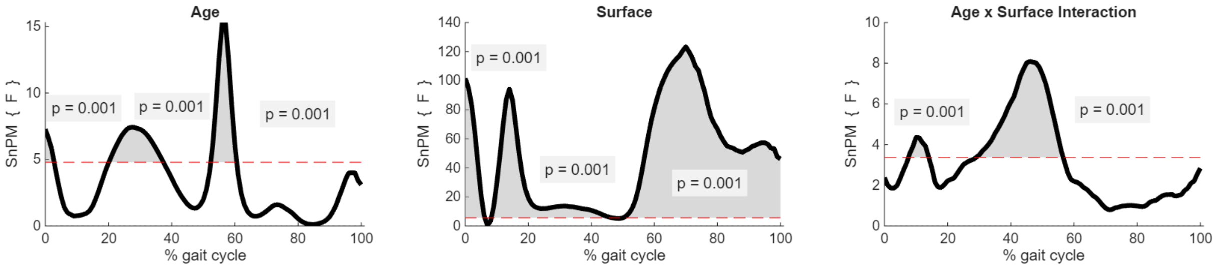


**Ankle-Knee DP**

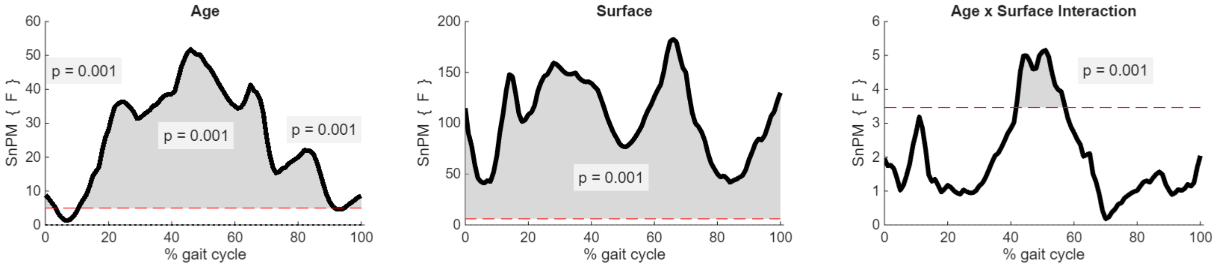


*Figure S2: ANOVA results for MARP and DP across the Knee-Hip and Ankle-Knee joint pairs*

### Mean joint kinematics

#### Hip joint

Surface comparisons

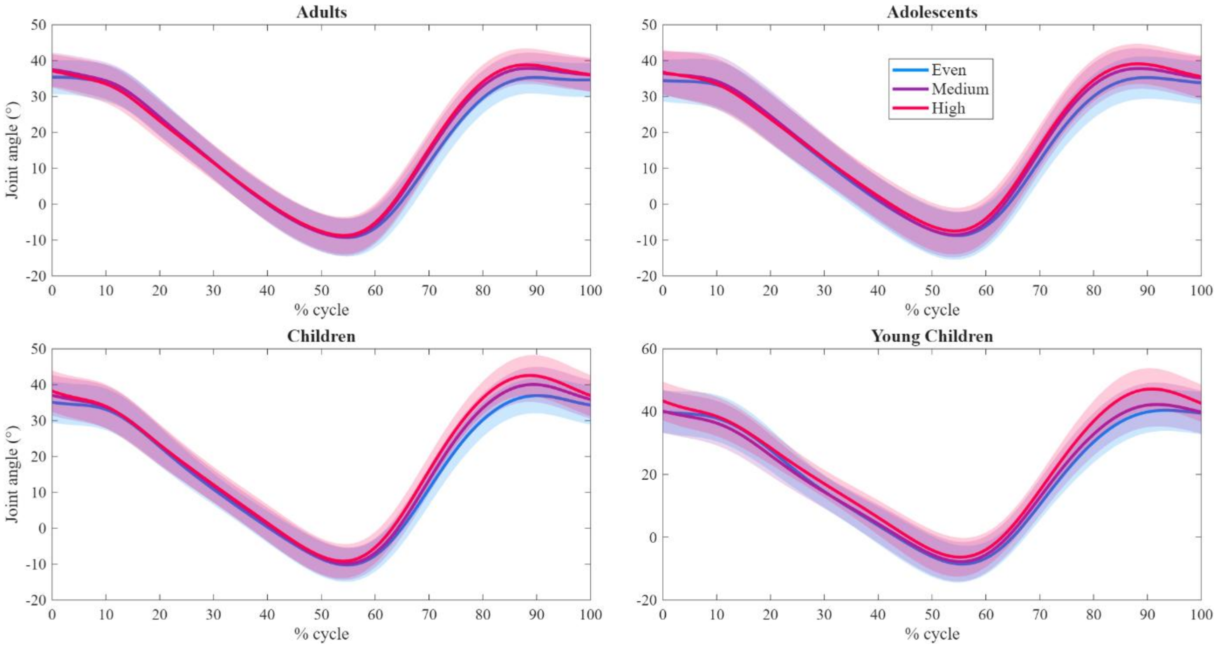


Age group comparisons

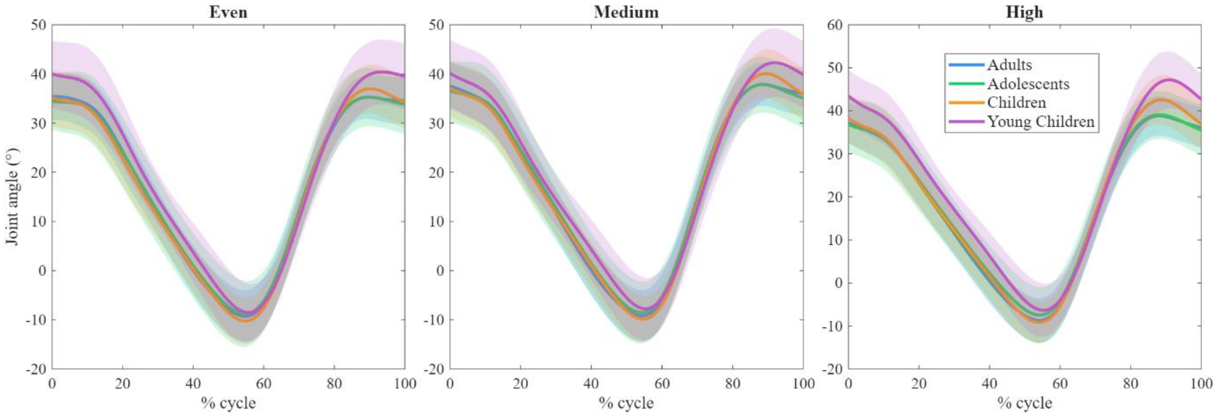


*Figure S3: Hip joint kinematics*

### Knee joint

#### Surface comparisons

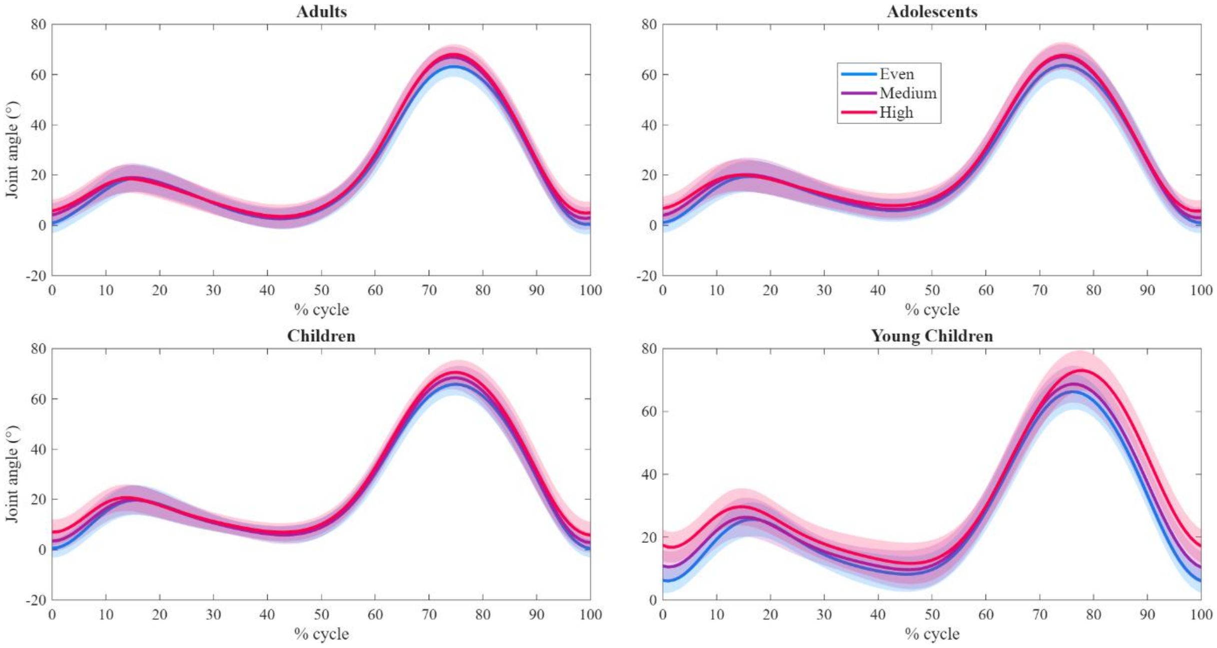


#### Age group comparisons

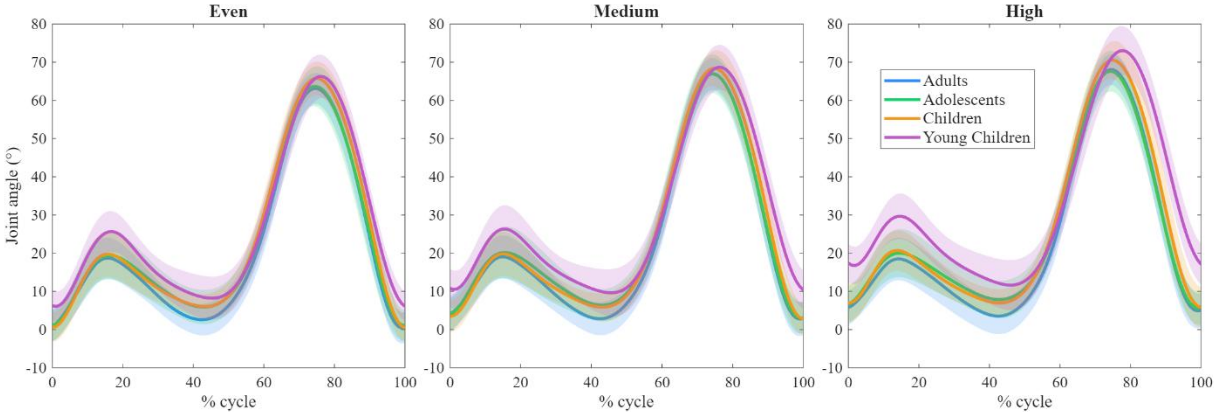


*Figure S4: Knee joint kinematics*

### Ankle joint

#### Surface comparisons

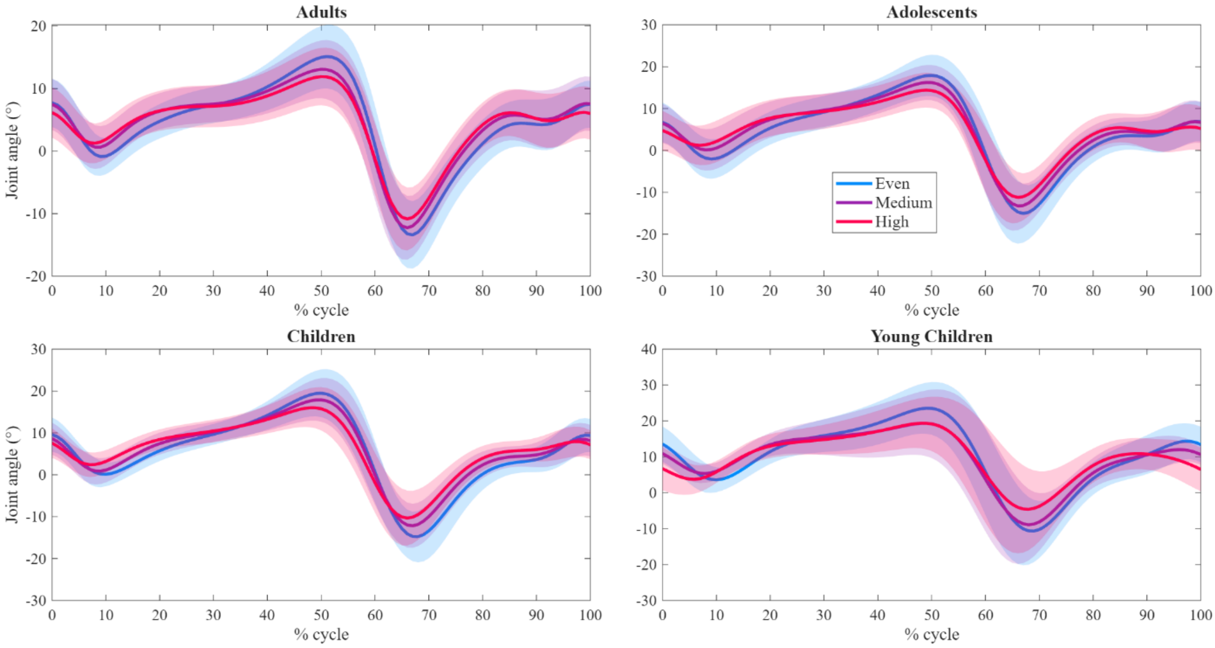


#### Age group comparisons

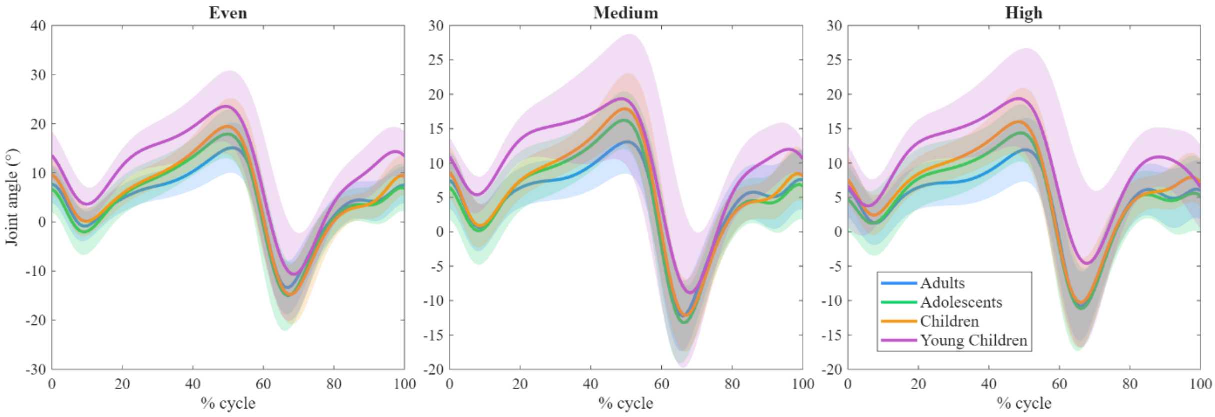


*Figure S5: Ankle joint kinematics*

### Inter-cycles joint kinematic variability

#### Hip joint

##### Surface comparisons

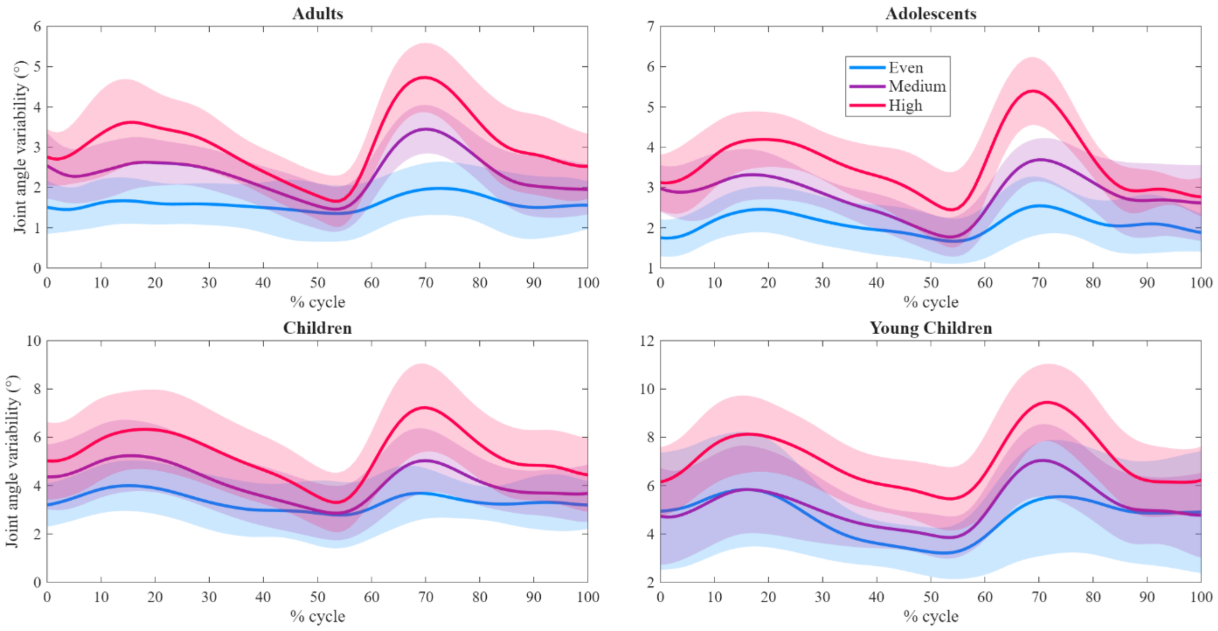


##### Age group comparisons

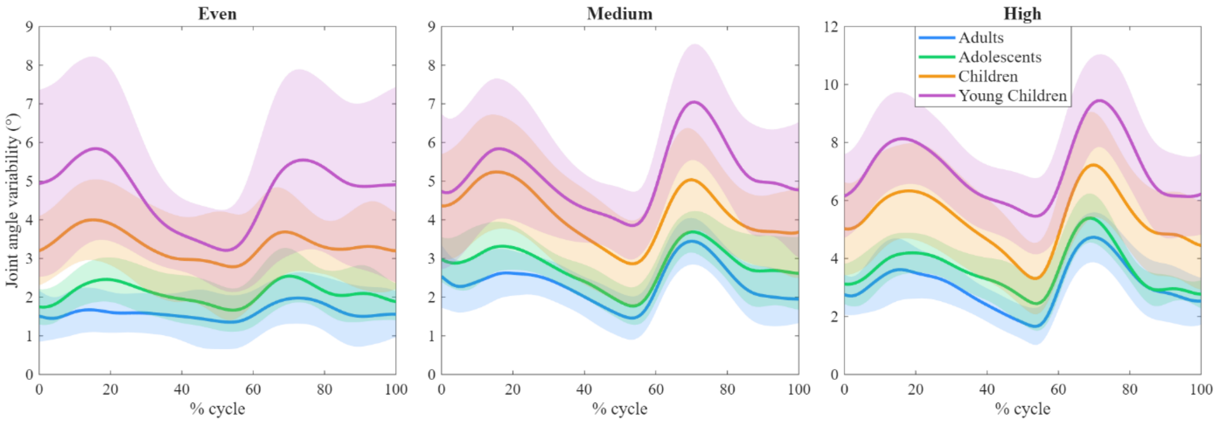


*Figure S6: Hip joint kinematic variability*

### Knee joint

#### Surface comparisons

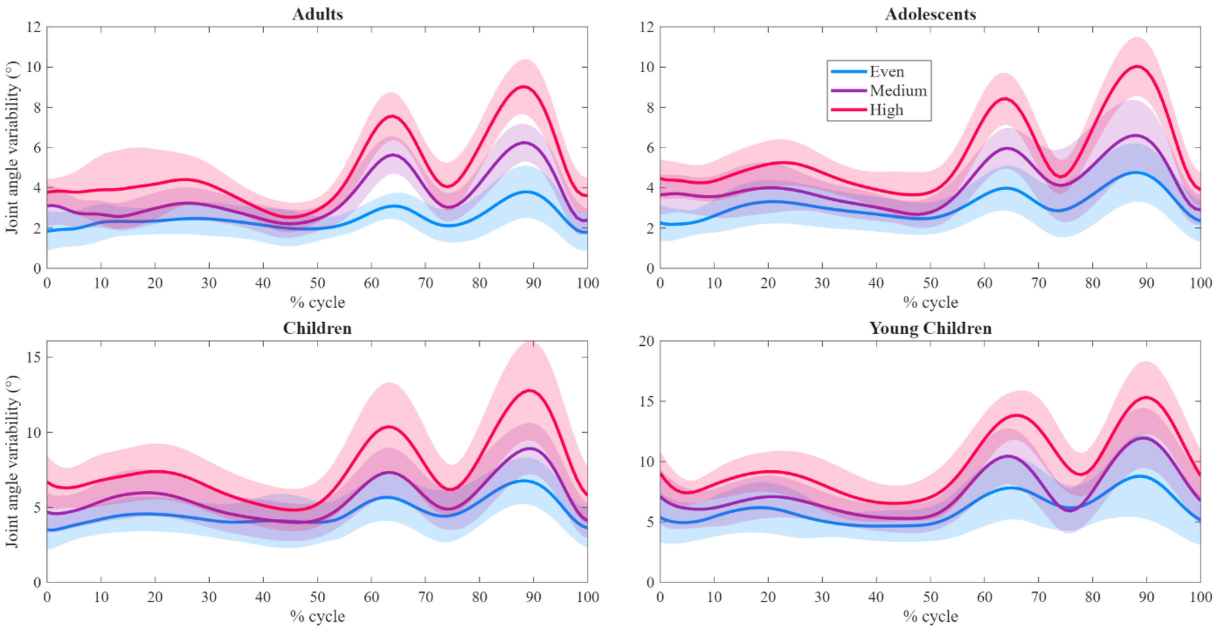


#### Age group comparisons

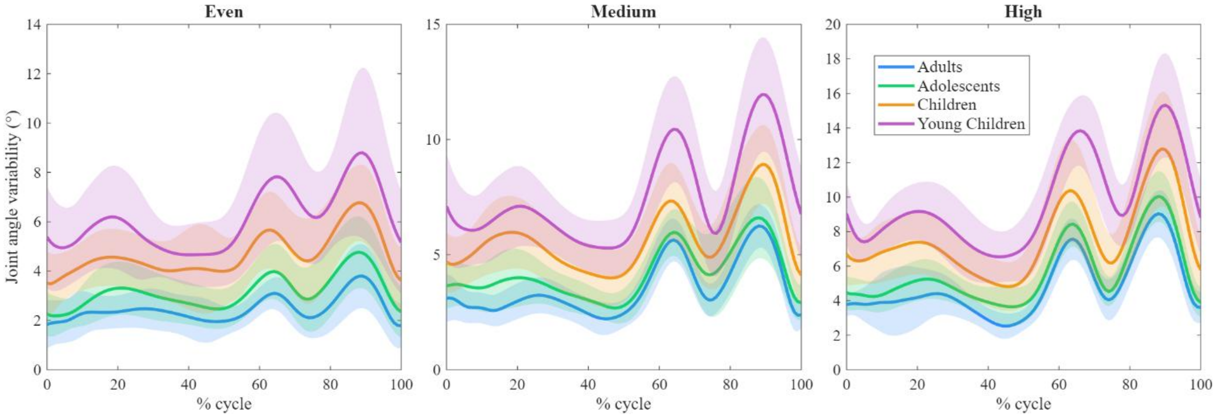


*Figure S7: Knee joint kinematic variability*

### Ankle joint

#### Surface comparisons

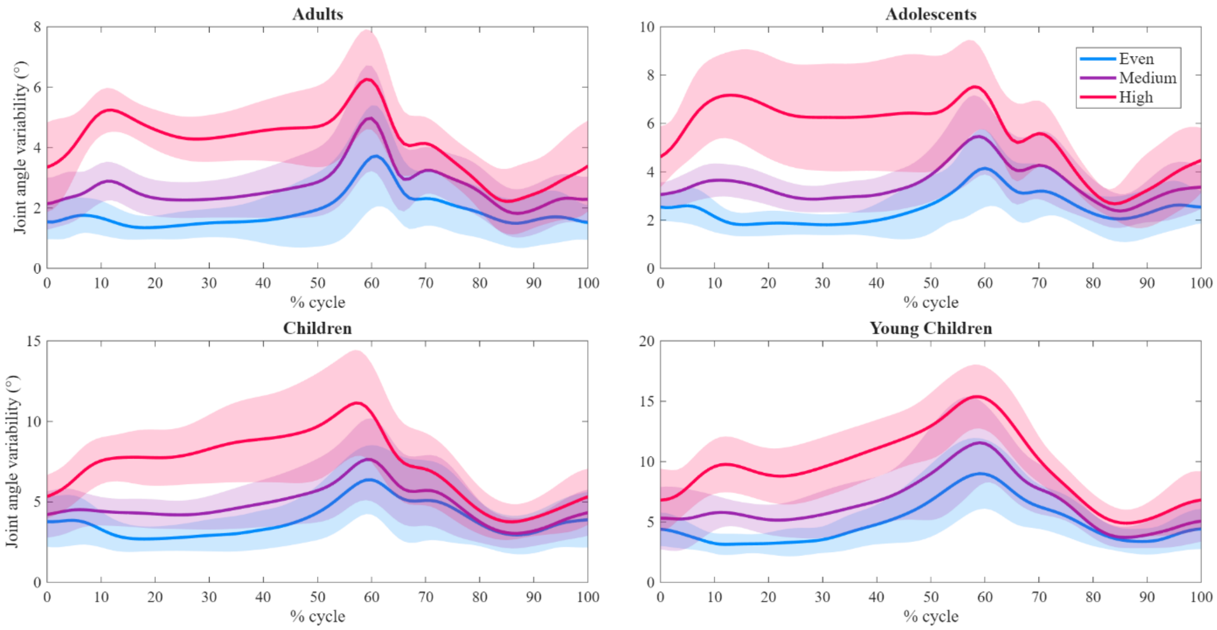


#### Age group comparisons

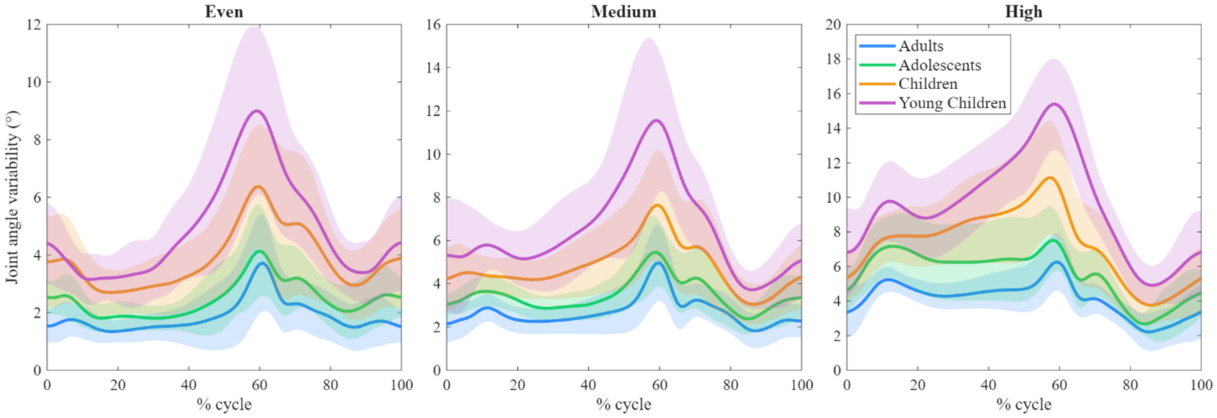


*Figure S8: Ankle joint kinematic variability*
